# An Axiom SNP genotyping array for potato: development, evaluation and applications

**DOI:** 10.1101/2025.08.17.670748

**Authors:** Nadia Baig, Kathrin Thelen, Mathieu A.T. Ayenan, Stefanie Hartje, Evelyn Obeng-Hinneh, Rafal Zgadzaj, Juliane Renner, Katja Muders, Bernd Truberg, Arne Rosen, Vanessa Prigge, Julien Bruckmüller, Jens Lübeck, Delphine van Inghelandt, Benjamin Stich

## Abstract

Potato is a versatile food crop and a major component of human nutrition worldwide. Genomic-assisted breeding methods have the potential to increase the gain of selection. We report the development and validation of a high-density Axiom-based SNP array for potato (*Solanum tuberosum* L.). Whole-genome 10X Genomics based sequencing of 108 diverse clones representing landraces, improved cultivars, and wild relatives identified around 23.8 million sequence variants, from which 929,127 variants, alongside with 18,718 markers from a previously developed Illumina Infinium 21K array (GGP3), were tiled on the array. The array demonstrated high reproducibility, with replicate samples showing an average concordance of 99.88% in genotype calls for PotatoTools specific variants, compared to 99.93% for the Illumina Infinium 21K array variants. A panel of 1,247 diploid and tetraploid clones was genotyped with the developed array. Genotype calling, considering allele dosage, was realized using fitpoly and yielded 852,793 calls. The array informativeness was optimized by applying Euclidean distance, heterozygous strength offset metrics, call rate, minor allele frequency filtering, and diploid-based allele correction, yielding a final set of 206,616 robust and informative markers. The filtered marker set enables precise characterization of genetic variation across diverse germplasm, thereby supporting robust analyses of population structure, genome-wide association studies (GWAS), and genomic prediction in potato. Population structure analysis genotyped clones revealed clear subpopulation differentiation consistent with known ploidy groups. In addition, our discriminant analysis of principal components revealed a weak but structured diversity among clones of different market segments. GWAS analysis of 998 potato clones identified sequence variants significantly associated with polyphenol oxidase (PPO) activity, confirming the platform’s efficacy for trait mapping. For the same trait, genomic prediction accuracies of 0.72-0.86 have been observed. The developed potato SNP array provides a robust platform for high-throughput genotyping, supporting genetic diversity studies, association mapping, and genomic-assisted breeding in this important crop.

## INTRODUCTION

The cultivated potato (*Solanum tuberosum* L.) is a widely grown tuber vegetable rich in protein, fibre, carbohydrate, and vitamins (Camire et al., 2009, Bradshaw and Ramsay, 2009, Reddy et al., 2018). It is the most important non-cereal crop in the world, after rice and wheat, and is consumed by over a billion people worldwide (Dolničar, 2021, Oppenheim et al., 2019, Birch et al., 2012). Especially in developing countries, the potato plays a significant role in providing nutritional security and is thus considered as a food for the future (Burgos et al., 2019, Thiele et al., 2010, Watanabe, 2015, Devaux et al., 2019).

Amidst the increasing impact of climate change, agriculture faces unprecedented hurdles, and potato production is no exception (Hijmans, 2003). Rising temperatures, unpredictable weather patterns, reductions of arable land or the surging population of pests pose significant threats to the potato crop production (Haverkort and Verhagen, 2008, Adavi et al., 2018, Quiroz et al., 2018, Handayani et al., 2019, Haq et al., 2023). Since production is becoming difficult, considerable yield increases are needed to meet rising demands for potato (Devaux et al., 2021). However, the genetic gain made in potatoes, especially in recent decades in terms of yield is lagging behind dramatically (Jansky, 2009, Lindhout et al., 2011, Stokstad, 2019) compared to other major crops such as maize, rice, and wheat (Fischer and Edmeades, 2010). The tetraploidy of cultivated potato poses a significant hurdle in potato breeding efforts resulting in a low gain of selection (Jansky, 2009) in complex traits such as yield. Moreover, a high degree of heterozygosity, long breeding cycles, and especially, the possibility of assessing complex traits such as yield only late in the breeding process are other reasons for low genetic gain of selection in potatoes (Hamilton et al., 2011, Xu et al., 2011, Hirsch et al., 2013, Spooner et al., 2014, Bradshaw, 2017, Stich and Van Inghelandt, 2018, Ortiz and Mihovilovich, 2019, Ghislain and Douches, 2019).

Genetic gain in potatoes can be increased by a thorough knowledge of the genetics of the traits under consideration (Bradshaw, 2022). The availability of well-established reference genomes (Van Os et al., 2006, Aversano et al., 2015, Leisner et al., 2018, Zhou et al., 2020, Freire et al., 2021, Wang et al., 2022, Hoopes et al., 2022, Sun et al., 2022) has opened doors for utilising molecular genetic tools in potato breeding. One way to improve genetic gain is to implement marker-assisted selection (MAS) (e.g. Naeem et al., 2021, Gebhardt, 2013, Pajerowska-Mukhtar et al., 2009). So far, targeted breeding approaches focused on identifying genes and molecular markers have enabled breeders to develop improved potato varieties with regards to qualitative traits (Angmo, D., Sharma, S.P. Kalia, 2023, Gebhardt et al., 2011). Moreover, SNP genotyping arrays such as the 96 BeadXpress SNP array (Hamilton et al., 2011), 8K Illumina Infinium array (Felcher et al., 2012), and 20K SolSTW array (Vos et al., 2015) have emerged as indispensable tools for potato breeders in the context of MAS for the characterisation of genome-wide diversity (Stich et al., 2013). Furthermore, they are essential as a powerful tool for genomic prediction that potentially allows increasing the gain of selection of potato for complex traits (Stich and Van Inghelandt, 2018, Endelman et al., 2018, Wu et al., 2023). The available SNP genotyping arrays have transformed potato breeding and research, yet they come with several existing limitations. These arrays have been developed using SNP discovery studies involving only few clones e.g. six varieties (Hamilton et al., 2011). The small size of the discovery panel introduces bias in identifying sequence variants, markedly missing rare variants compared to the common variants due to shifts in allele frequencies. In addition, all available arrays lack comprehensive genome coverage (Moragues et al., 2010), limiting their suitability for gene discovery projects and for genomic prediction, depending on the diversity of the genetic material (Marhadour and Prodhomme, 2022).

An alternative to array-based genotyping is exploiting the possibilities that have arisen from the considerable reductions in sequencing costs (Poland and Rife, 2012, He et al., 2014) such as genotyping by sequencing (GBS). GBS offers comprehensive genome-wide coverage and is particularly valuable for discovering novel genetic variants (Elshire et al., 2011, Poland and Rife, 2012). The higher computational complexity for obtaining genotype calls compared to SNP array data makes GBS less appealing for routine genotyping in the context of breeding programmes. However, the even more important point is that obtaining dosage information at each sequence variant requires the coverage in GBS studies of polyploid species, such as potato, to be increased considerably compared to diploids, which increases the cost per data point (Gemenet et al., 2020). Therefore, potato breeding programmes prefer high-density SNP genotyping arrays due to their cost-effectiveness, speed, and precision in capturing high degrees of genetic information.

In the context of these challenges and opportunities, this study aims to (i) describe the design of a high-density Axiom based SNP genotyping array for potatoes, (ii) assess the effectiveness of the array and, (iii) present the utility of this array to infer genetic diversity and population structure of a diverse set of potato clones and its application for genome-wide association mapping and genomic prediction.

## MATERIALS AND METHODS

### Plant materials, library preparation and sequencing

The resequencing panel used in this study consisted of a total of 108 clones (Table S1). Of these, 7 were non-tuberosum diploid, 1 non-tuberosum tetraploid which together have hereafter been designated as PotatoTools NonCult21PT 950K clones. The other 100 were tetraploid cultivated potato clones referred to as PotatoTools Cult21PT 950K clones.

The clones were selected to represent all market segments as well as important cultivation areas (Europe, East Asia, North and South America) (see Table S1) fulfilling one or several of the following criteria: (i) cultivars with high significance in breeding; (ii) promising breeding clones; (iii) parental genotypes of frequently used mapping populations; (iv) wild species which have significance as disease resistance donors. All maturity groups are of very early to mid-late maturity.

High molecular weight genomic DNA was extracted from frozen leaf material following Mayjonade et al. (2017). For each sample, paired-end libraries were prepared using the Genome reagent kit v2 from 10x Genomics following the recommended protocol. The DNA libraries were sequenced on Illumina Hiseq 2500 and NovaSeq S4 platforms aiming for 2×150 bp reads. A customised bioinformatics workflow was designed to process 50.94 billion raw reads, followed by sequence variant identification, quality filtering, and sequence variant selection (Figure S1).

### Data pre-processing and variant identification

Raw reads of the 108 clones were mapped to the potato reference genome of a diploid clone derived from the elite cultivar Agria (Freire et al., 2021) using longranger’s *align* function with default settings (https://github.com/10XGenomics/longranger) as this genome offered the highest mapping rates at the time of analysis. Local realignment around indels was performed using Genome Analysis Toolkit (GATK-version 3.7) (McKenna et al., 2010). PCR duplicates were removed using Picard (v.1.130) and uniquely aligned paired reads having a mapQ score of ≥ 20 were retained for variant calling (Hardigan et al., 2017).

The BAM files of were divided into two groups based on their germplasm status: PotatoTools i) Cult21PT 950K and ii) NonCult21PT 950K clones. The filtered BAM files were processed using Freebayes to make combined sequence variant calls for group (i) and individual calls for clones of group (ii) following Garrison and Marth (2012). The aim was to group all cultivar clones and get sequence variants specific to important non-tuberosum clones with significance as disease-resistance donors before applying custom parameters for sequence variant filtering and selection.

### Sequence variant filtering

Quality filtering of bi-allelic sequence variants was performed using a custom Python script in combination with Vcflib (v1.0.2) and bcftools (v3.6.5) to keep sequence variants with a (i) Phred-quality score ≥ 20; (ii) average read depth ≥15; (iii) depth (DP) lower than the average sequence DP divided by the total number of samples plus three standard deviations; (iv) missing rate per sequence variant ≤20% and (v) minor allele frequency (MAF) *>* 0.05 (Hardigan et al., 2017, Vos et al., 2015). Sequence variants unique to each clone of PotatoTools NonCult21PT 950K were obtained using bcftools (v3.6.5). The goal was to retain alleles unique to individual clones of NonCult21PT 950K while excluding any shared alleles between them, thus ensuring the unique genetic variation of the clones.

### Sequence variant annotation and initial selection for *p convert* value calculation

Amino acid substitutions and their impact on protein function were estimated using SIFT4G (v2.4) software (Vaser et al., 2016) based on the Agria reference genome (for details see Method S1). A customized sequence variant selection pipeline (Figure 1) was designed to select a subset of 32,000 sequence variants from PotatoTools NonCult21PT 950K clones and 2.85 million from PotatoTools Cult21PT 950K clones following Affymetrix guidelines (http://www.affymetrix.com/) for which the Affymetrix *p convert* value was calculated. This customised pipeline (see details Method S2) ensured the selection of sequence variants that are uniformly distributed across the genome by using a custom genome binning approach. From each bin, the sequence variants having no interfering variant nearby were prioritised compared to the ones having one or more interfering variants for selection as they are likely to be successfully converted to an informative probe on the array. The A/T, C/G variants and indels were de-prioritised as they require more space on the array. Additionally, the variants exhibiting higher MAFs were prioritised for selection and scoring. For the PotatoTools NonCult21PT 950K clones specific variants, a binning criterion was again followed, where the equal representation of both intergenic and genic mutations was ensured. Lastly, the selected variants plus the 21,227 sequence variants from the Illumina Infinium 21K array (GGP3) were submitted to Affymetrix for *p convert* score evaluation. The *p convert* score categorised sequence variants as “recommended”, “neutral”, “not recommended” and “not possible”. Sequence variants tagged as “recommended”,“not recommended” and, “neutral” were further analysed for final array tiling by reapplying the variant selection based on a binning approach which is described in the next section.

**Figure 1:**
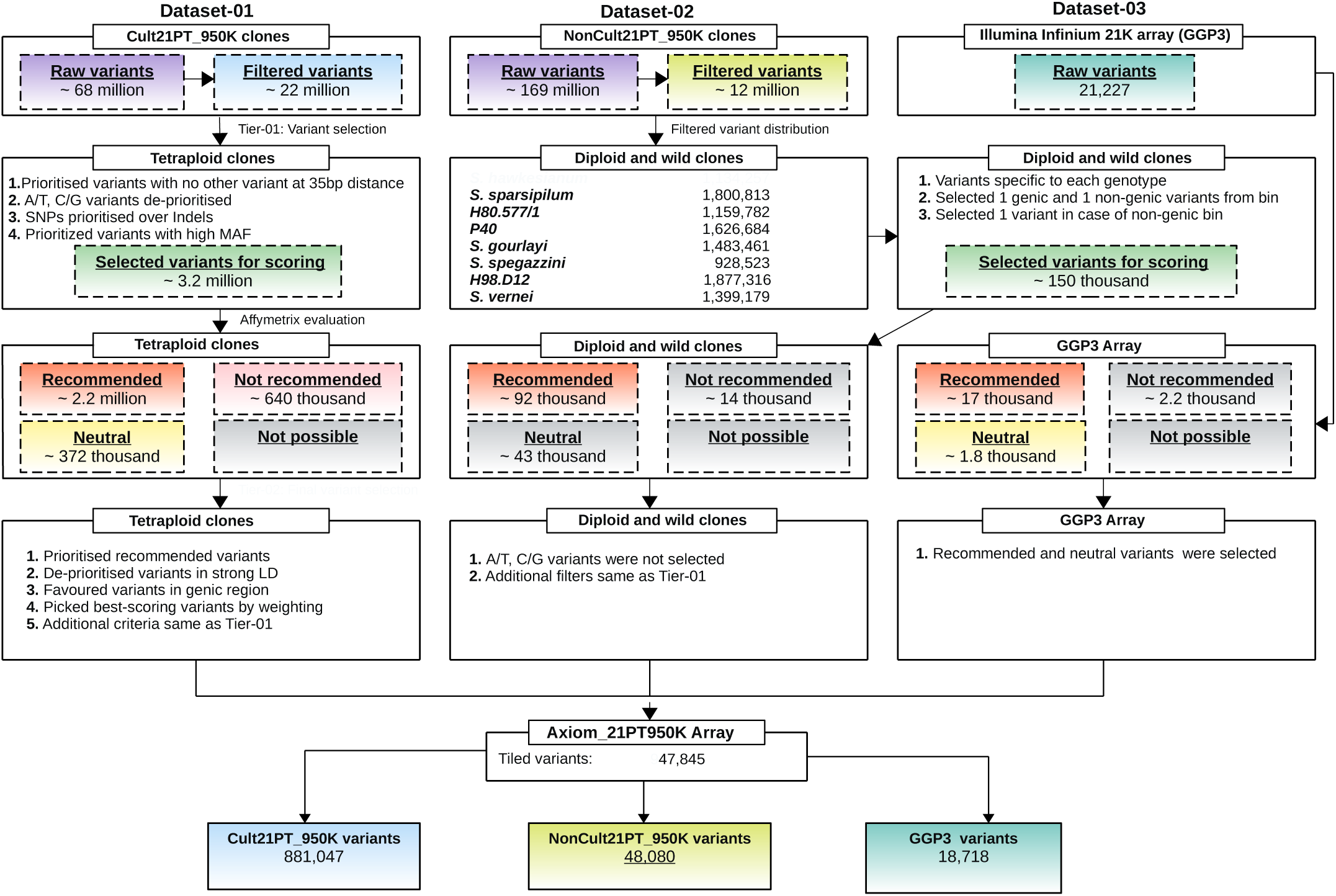
Workflow with the major sequence variant filtering and selection steps for the design of the Axiom_21PT_950K screening arrays. Workflow illustrating filtering and sequence variant selection steps for PotatoTools Cult21PT_950K, NonCult21PT_950K clones, and Illumina Infinium 21K array (GGP3) for the development of Axiom 21PT 950K screening arrays for potato.

### Sequence variant selection for the development of two Potato Axiom 21PT 950K screening arrays and linkage disequilibrium analysis

Final variant selection for inclusion on the Axiom 21PT 950K screening arrays was performed by adopting a similar strategy as above to distribute 50% of the sequence variants uniformly in each 5kb segment. The other 50% of the sequence variants were distributed according to the gene density per window providing a higher number of sequence variants in gene-rich parts of the genome. The balance between a uniform distribution of variants relative to the gene density, as well as the selection criteria including MAF, A/T and C/G variant types, linkage disequilibrium (LD), proportion of missing genotypes at each polymorphism, variant effect and *p convert* value, was optimised using the following selection index approach: (i) Sequence variants obtained after *p convert* calculation were divided into four groups based on their MAF and assigned a score of 1 to 4. A score of 1 was assigned to the group of sequence variants having 0.05 ≤ MAF*<* 0.15, 2 to the variants exhibiting 0.15 ≤ MAF*<* 0.25, and a score of 3 and 4 was assigned to variants having 0.25 ≤ MAF*<* 0.35, and MAF≥ 0.35, respectively. (ii) All A/T and C/G sequence variants were assigned a score of 1 as these take up more space on the array, while all other variants obtained a score of 2. iii) Four categories of sequence variants were made based on the average extent of linkage disequilibrium (LD) measured as *r*^2^ between the target sequence variant and the neighbouring variants within 25kb up and downstream. Sequence variants having 0 ≤ *r*^2^ ≤ 0.15 were assigned a score of 4, sequence variants with 0.15 *< r*^2^ ≤ 0.3 were assigned a score of 3, sequence variants having 0.3 *< r*^2^ ≤ 0.5 and 0.5 *< r*^2^ ≤ 1 were assigned scores of 2 and 1, respectively. (iv) Sequence variants designated as “recommended” according to their *p convert* value (*p convert >* 0.6) (Montanari et al., 2019) were assigned a score of 3, “neutral” were assigned a score of 2, and “not recommended” and “not possible” variants were not considered. (v) Indels were de-prioritised by assigning them a score of 1, while SNPs were assigned a score of 2 as indels require complex probe designs for accurate detection and, thus, more space on the array. (vi) Based on the SIFT4G annotations, sequence variants present in genic regions were scored 4, and sequence variants in intergenic regions as 3. For the final index value calculation, the score of sequence variant attributes (i), (v) and (vi) were multiplied with a weight of 3, while the score for the rest of the attributes was multiplied by 2, to ensure the high performance of the final screening array. (vii) The sequence variants from Illumina Infinium 21K array (GGP3) were prioritised over PotatoTools Cult21PT 950K and NonCult21PT 950K clones specific variants by assigning them a score of 200. (viii) Sequence variants having no missing genotype information in all clones were given a score of 2 and those having up to 20% missingness were given a score of 1. For the final index value calculation, the scores of sequence variant attributes (i), (v) and (vi) were multiplied by a weight of 3, while the score for the rest of the attributes was multiplied by 2. The index value for each sequence variant was subsequently calculated by aggregating the scores of the respective attributes described above.

### Genotyping and array performance evaluation

Genomic DNA was isolated from the validation set with 1,247 potato clones including the 108 clones from the above-mentioned resequencing panel. The additional 1,139 potato clones were selected from the first micro plot stage (A clone) from the potato breeding programmes of Saka (Saka Pflanzenzucht GmbH & Co. KG), Norika (Nordring-Katofellzuchtund Vermehrungs-GmbH) and EUROPLANT Innovation GmbH Co. KG and were designated below as PotatoTools populations. The entries of the PotatoTools populations belong to 171 full-sib families. The number of clones within a population varied from 1 to 38 clones. The parental clones of these segregating populations were included as well.

The selected clones belonged to four different market segments: table potato, crisps, french fries and starch processing/industrial use. One control sample was placed on each of the thirteen, 96-well plates as a technical replicate. The cell intensity files from the Axiom assay were subjected to quality control using the Axiom Analysis suite according to best practice. The dish quality filter DQC ≥ 0.82 was applied to the 1,247 clones. To include all PotatoTools NonCult21PT 950K clones, the QC threshold for this group was relaxed and all clones exhibiting a QC call rate ≥ 78% were retained for the downstream analyses.

Marker dosage was estimated from ratios of signal intensities of the Potato Axiom 21PT 950K screening arrays using the R package fitPoly (v.3.0.0). The default settings of fitPoly were adjusted: (i) allowing 95% confidence for a genotype to belong to a specific cluster (p.threshold = 0.95), (ii) peak threshold value was set to 0.99 to allow sequence variants with low allele frequencies to be fitted, (iii) call.threshold was set to the default value of 0.60, allowing the rejection of markers with more than 40% missing values and (iv) ploidy was set to 4 and diploid clones were analysed together with tetraploids to allow the confirmation of correct nulliplex (AAAA), duplex (AABB) and quadruplex (BBBB) clusters. The R package SNPolisher (v.1.5.2) was used in the next step to visualize the cluster plots of fitted sequence variants. The fitted calls were additionally filtered by applying a SNP call rate (CR) threshold of ≥ 90%. This threshold is defined as the ratio of the number of samples assigned a genotype call other than “NoCall” to the total number of samples. Additionally, only sequence variants having MAF of ≥ 0.01 were included in the downstream analyses.

We evaluated the informativeness and performance of the Potato Axiom 21PT 950K screening arrays through an investigation of (i) genotyping call reproducibility of Illumina Infinium 21K array (GGP3) vs. Potato Axiom 21PT 950K screening arrays, and (ii) technical replicates-based reproducibility for the filtered sequence variants.

### Selection of highly informative sequence variants for genotyping

We used exceptionally stringent filtering criteria aiming to select the most reliable sequence variants. For the final selection of the sequence variants to be included on a Potato Axiom production array, we calculated the minimum Euclidean distance between the centres of the heterozygous clusters and those of the homozygous clusters in the X dimension (AB) (Figure 2) by considering sequence variant posterior values. Furthermore, heterozygous strength offset (HetSO) metrics, which measure how far the heterozygous cluster centre sits above or below the centres of the homozygous cluster in the Y dimension (*A* + *B*)*/*2 (Figure 2) was calculated using a customised Python script.

**Figure 2:**
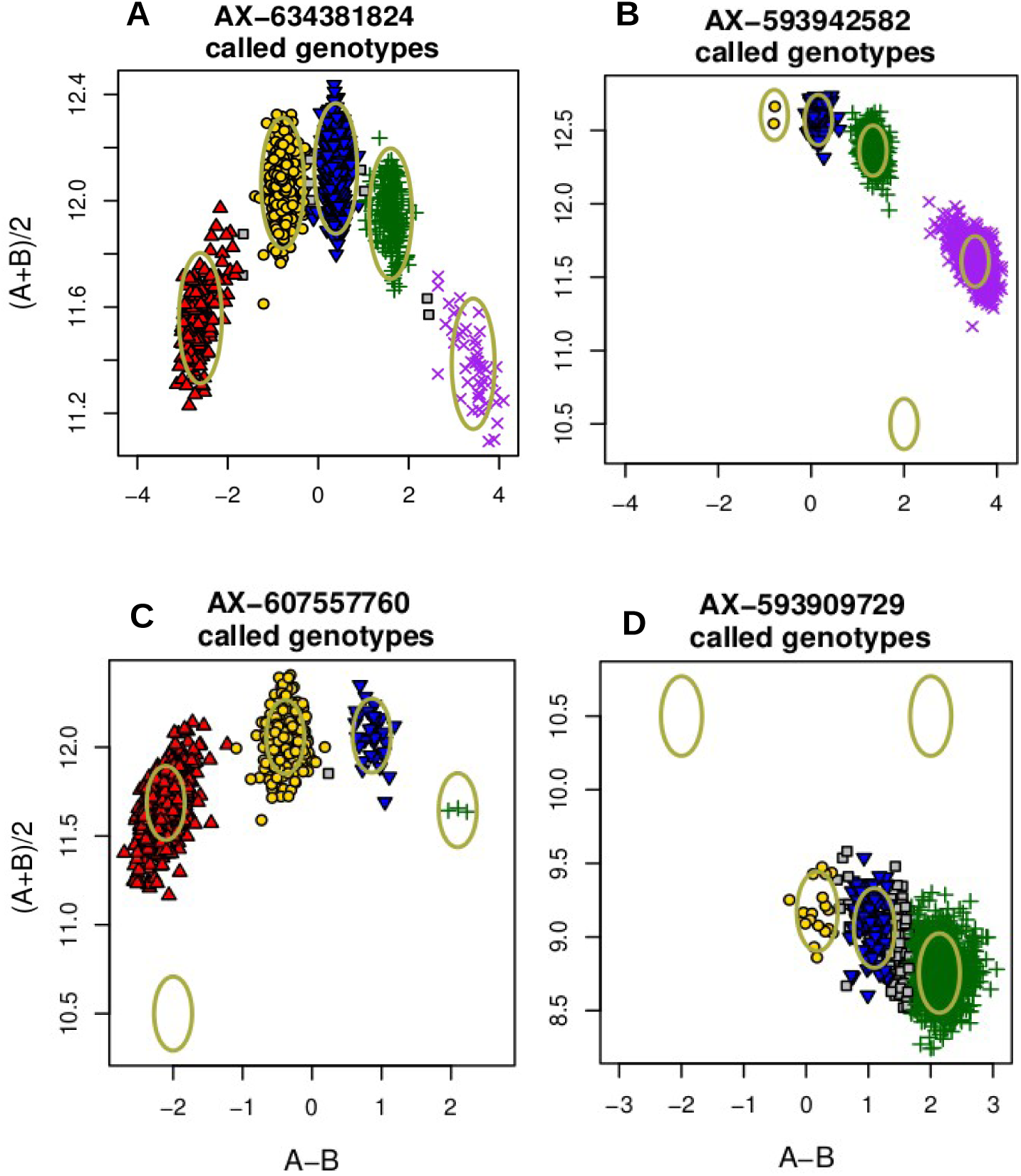
Sequence variant classification based on the distribution of genotypes into five genotype clusters: red points indicate AAAA, yellow points indicate AAAB, blue points represent AABB, green points indicates ABBB, and purple points indicate BBBB clusters for example sequence variants. (A) Cluster Type-I sequence variants with both homozygous and all heterozygous clusters present, (B) Cluster Type-II sequence variants having the AAAA homozygous cluster missing, (C) Cluster Type-III sequence variants with a missing homozygous cluster BBBB, and (D) Cluster Type-IV sequence variants with both homozygous clusters missing.

The sequence variants were classified into five classes based on the distribution of genotypes carrying A (reference) and B (alternate) alleles in the clusters. Sequence variants with AAAA, BBBB, AABB, ABBB and AAAB clusters ≠ 0, where at least one clone is present in each genotype cluster, were labelled as Cluster Type-I, clusters where AAAA==0 but the rest ≠ 0 were grouped into Cluster Type-II, BBBB==0 with the rest ≠ 0 being grouped into Cluster Type-III, AAAA & BBBB==0 while the rest ≠ 0 clusters were grouped into Cluster Type-IV, and all other combinations of missing genotype clusters were grouped into Cluster Type-V. We filtered sequence variants for an Euclidean distance ≥ 0.05 and a HetSO ≥ -0.1. These stringent criteria were applied to identify variants with well-separated cluster centres.

In later steps, sequence variants were further inspected to identify variants with incorrect genotyping results by examining if genotype calls for diploids were made for the simplex (AAAB) or triplex (ABBB) clusters. Sequence variants exhibiting wrongly assigned calls for diploid clones where empty homozygous clusters (Cluster Type-II, Cluster Type-III, and Cluster Type-IV) were observed were subjected to leftwards and rightwards genotype shifting to evaluate whether correction of the diploid genotype is possible: In the case of the missing AAAA cluster (Figure 2B), all available genotype calls were shifted one cluster leftwards. For the sequence variants with no genotype calls in the BBBB cluster (Figure 2C), all available genotype calls were shifted rightwards. For cases with no genotype calls in both homozygous clusters (Figure 2D), we evaluated the shifting of all genotype calls in both directions. We kept for all sequence variants those shifted genotypes that solved all the genotype calls for diploids that were incorrect (i.e. ABBB or AAAB) in the previous calling. We removed all the sequence variants where allelic shifting did not correct the wrongly assigned calls for diploids. The filtered sequence variants alongside the genotype-corrected calls were then used for the downstream analyses.

### Population structure and clustering

Population structure was assessed for all potato clones that were part of the resequencing panel and the PotatoTools populations, as well as their parental clones by performing a Principal Component analysis (PCA) and a Discriminant analysis of principal components (DAPC). The DAPC analysis was performed using the “dapc” function of adegenet (Jombart et al., 2010) and the “find.clusters” function. The find.clusters function was used to identify the genetic clusters using K-means clustering. Overall, 100 principal components (PCs) were retained for dimensionality reduction. The number of clusters was chosen as five based on the Bayesian Information Criterion (BIC). We performed cross-validation using the “xvalDapc” function with 30 replicates and 90% of the individuals in the training set to prevent overfitting and to optimize the number of principal components retained in the DAPC. A total of 80 PCs minimising the Mean Squared Error (MSE) were selected as the optimal input for DAPC and the first four discriminant functions.

### Genome-wide association mapping (GWAS) and genomic prediction analysis

#### Field experiments and phenotyping

The PotatoTools populations but not their parental genotypes were phenotypically characterised in field experiments. The phenotypic data were collected in 2019, 2020, and 2021. The data from 2019 was collected from one location for each company (EUROPLANT: Kaltenberg, Norika: Gross Lüsewitz, and Saka: Windeby). For 2020 and 2021, a second location (EUROPLANT: Böhlendorf, Norika: Mehringen, Saka: Gransebieth) was selected by each breeding company resulting in a total of five different environments per company. Overall, there were a total of 15 environments. The clones were organised in blocks and each block comprised rows and columns. The number of plants per plot ranged from 9 to 20 depending on the corresponding environment (see Table S9). Eight “check” varieties were grown by each breeding company in all blocks of all environments, where the entries were cultivated by only one company for ownership reasons. For 2021, the experiments from Saka were performed in two trials separating the clones by maturity. One trial had clones from extra early to early maturity and the second trial included material of middle early to mid late maturity. Overall, this experiment comprised 5,947 observed plots. Phenotypic data was collected for the trait PPO, which is the enzymatic activity of polyphenol oxidase causing the browning of potato tubers. Potato breeders focus on reducing PPO activity to minimize enzymatic browning, which enhances product appearance, improves bruising resistance and extends shelf life, ensuring better quality for consumers and the processing industry. Since PPO activity is correlated with bruising scores, many breeders use it as a secondary trait in selection for bruise-resistant varieties. The trait values were evaluated by assigning a rating from 1 to 9 based on the browning of the potato tuber discs after the application of mechanical impact on tubers in a shaking device (cf. Urbany et al., 2011). This data was used for both the GWAS and genomic prediction analysis explained below. The phenotypic data from 15 environments were analysed according to the following model:

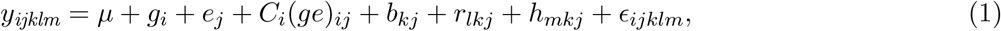

where *y_ijklm_* is the phenotypic observation of the *i^th^* potato clone in the *m^th^* column and the *l^th^*row of the *k^th^*block in the *j^th^*environment. The *µ* is the intercept term, *g_i_* the effect of *i^th^* clone, *e_j_* the effect of *j^th^* environment, *C_i_* the dummy variable filtering for checks with *C_i_* =1 and *C_i_*=0 for entries, and (*ge*)*_ij_* the interaction effect of the *i^th^* clone and the *j^th^* environment, *b_kj_* is the effect of *k^th^* block of the *j^th^* environment, *r_lkj_* the effect of *l^th^*row of the *k^th^*block of the *j^th^*environment, *h_mkj_*the effect of *m^th^*column of the *k^th^*block of the *j^th^*environment, and *c_ijklm_* the residual error. All effects were considered random except for *g_i_*. The records with standardised residual values greater than 3.5 or smaller than -3.5 were considered outliers and were removed from the analysis after visual inspection. In the next step, the adjusted entry means for all clones (checks and entries) were calculated across all environments according to equation (2).

In this step, residual error variance was assumed to be heterogeneous for the different environments.

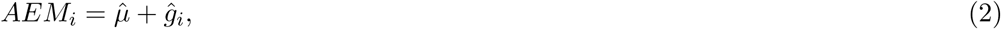

where, *µ*^ is the estimate of the intercept and *g*^*_i_* the estimate of the *i^th^* clone. In the calculation of the genetic variance, the clone effect was divided between the checks and the entries, where we used the following model with the effect of checks considered fixed, while the effect of the entries was considered random:

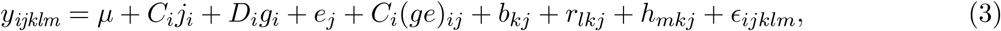

where *D_i_* is the indicator variable filtering for the entries with *D_i_* =0 for checks and *D_i_*=1 for entries. Furthermore, heritability on an entry mean basis was calculated for each trait according to the following equation:

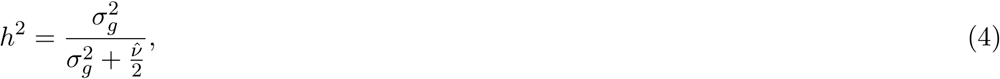

where *σ*_*g*_^2^ is the genotypic variance from model (3) and *v*^ denotes the mean variance of a difference of two adjusted treatment means of the entries as these were assessed by model (1) with heterogeneous error variances.

A total of 988 out of 1,066 clones used in the field experiment had both genotypic and phenotypic data and were included in the genome-wide association and genomic prediction analyses. Overall, 202,008 sequence variants fulfilled the following criteria: (i) biallelic loci, (ii) loci with *<* 20% missingness, and (iii) and loci with MAF ≥ 0.05 were used in the analyses. The missing sequence variants were mean imputed.

### Genome-wide association mapping

For the GWAS analysis, the above-mentioned adjusted entry means were used. A combination of SNP-based genomic relationship matrix (**G**-matrix) proposed by VanRaden (2008) and pedigree based **A**-matrix combined together as an **H**-matrix using the method described by Muñoz et al. (2014) was used. Sequence variants were considered fixed effects in the analysis using the following model:

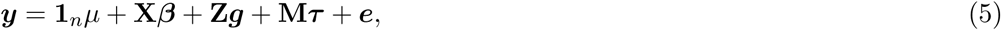

where ***y*** is the vector of adjusted means, **1*_n_*** the vector of ones, ***µ*** the overall mean and, ***β*** the vector of fixed effects that includes the first two principal components (P) of the sequence variants matrix to account for the population structure and X is the associated design matrix. g is the vector of random effect of the genetic background of each clone with variance = **H***σ*^2^ where ***H*** is the above mentioned genetic relationship matrix and ***Z*** is the associated design matrix. ***τ*** is the fixed effect of the sequence variants under consideration, ***M*** is the associated design matrix, ***e*** represents the vector of the residual variance and it is assumed to be normally distributed. Sequence variants having a significant association with PPO were determined based on the Bonferoni correction for multiple testing.

### Genomic prediction

The genomic best linear unbiased prediction (GBLUP) model was used to predict genomic estimated breeding values (GEBVs) for the PPO trait. The model can be represented as:

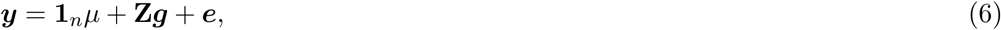

where ***y*** is the vector of phenotypic adjusted entry means calculated using model (3), **1*_n_*** the vector of ones, *µ* the overall mean, ***Z*** a design matrix mapping the adjusted entry means to clones, ***g*** the vector of genomic breeding values, and *e* the error term. The genomic breeding values ’***g***’ were considered to follow ***g*** ∼ N(0*, **G**σ*^2^) where *G* represents the above mentioned genomic relationship matrix considering Ashraf et al. (2016) an additive autotetraploid model.

The GBLUP model was calculated using the sommer package in R (Covarrubias-Pazaran, 2016). A training set (TS) was used to estimate the effects of the model and then used to calculate the genomic estimated breeding values (GEBVs) of the genotypes within the validation set (VS). A five-fold cross-validation approach (CV) was implemented in the predictions and the data was split randomly into five parts, where one part was used as VS and the rest as TS. Moreover, each of the five parts was used as VS once. This procedure was repeated 50 times. The model performance for PPO trait was measured by the prediction ability i.e. the Pearson correlation coefficient between the adjusted entry means (AEMs) from model (1) and GEBVs of the VS. Lastly, prediction accuracy (PA) was calculated as the ratio of prediction ability and the square root of heritability *h*^2^.

## RESULTS

### Sequence variant identification and array design

Approximately 5.1 TB of whole genome sequencing data comprising 50.95 billion mapped reads in PotatoTools Cult21PT 950K and NonCult21PT 950K clones were analysed in the current study for the identification of sequence variants to design two high-quality Potato Axiom 21PT 950K screening arrays. Using a customised bioinformatics workflow (Figure S1 and Figure 1) resulted in the detection of 68,317,505 raw sequence variants in the PotatoTools Cult21PT 950K (Table S2) and 169,989,471 in PotatoTools NonCult21PT 950K clones (Table S3). After stringent sequence variant selection by considering average read depth, missing rate per polymorphism, MAF, and Phred-quality score criteria (Figure S1), 67% (22.05 million) sequence variants were retained for the PotatoTools Cult21PT 950K clones (Table S2). For PotatoTools NonCult21PT 950K, on average 11.4 million sequence variants with private alleles were obtained after filtering (Table S3).

In our sequence variant selection workflow, we paid attention to the genome-wide distribution of sequence variants but also considered heterogeneity in gene density across the genome when selecting sequence variants. Adhering to Affymetrix-based filtering recommendations, we selected 3.2 million sequence variants from the PotatoTools Cult21PT 950K and 150,000 from the PotatoTools NonCult21PT 950K clones for predicting their conversion suitability ( i.e. *p convert* score). An additional set of 21,227 sequence variants from the Illumina Infinium 21K array (GGP3) were also scored to check their conversion suitability. Based on this evaluation, we selected a subset of 947,845 sequence variants, consisting of 881,047 variants from PotatoTools Cult21PT 950K, 48,080 from PotatoTools NonCult21PT 950K clones and 18,718 Illumina Infinium 21K array (GGP3) specific variants for tiling on the Axiom 21PT 950K screening arrays (Figure 1).

### Genome-wide distribution of the Potato Axiom 21PT 950K array sequence variants

The genome-wide analysis revealed a comprehensive distribution of sequence variants across the potato genome. The twelve chromosomes were densely covered by a total of 876,528 sequence variants, where the number of sequence variants within 1Mb windows ranged from 0 to 2200, resulting in an average spacing of 856 bp (median: 377 and standard deviation (*σ*) of 1,444). Quantitative analysis of the sequence variant distances revealed that 75.2% of variants have an adjacent sequence variant within less than 1kb and 18.9% have adjacent sequence variants within 1-3kb indicating a good coverage of the genome (Figure S2(A)). Moreover, on average, 12 sequence variants were found within 1kb of the coding sequence with *σ* of 8.7 variants in 1kb.

Furthermore, our analysis revealed that 21.7% of the sequence variants were located within 1kb of a gene (Table S4A). In contrast, Illumina Infinium 21K array (GGP3) sequence variants had an average spacing of 39,283 bp (median: 457 bp) due to lower variant density with an average standard deviation of 137,918 bp. Around 56.1% of the Illumina Infinium 21K array (GGP3) sequence variants had an adjacent sequence variant within 1kb, while 30.2% have an adjacent sequence variant at a distance ≥ 10kb (Figure S2(B)). The SIFT4G algorithm predicted amino acid substitutions affecting protein structure in 218,198 out of 929,127 tiled sequence variants of the Potato Axiom 21PT 950K screening arrays. The rest were unannotated due to insufficient data for accurate prediction. Scaffold chromosome variants were excluded from the SIFT4G based analyses. The highest percentage of variants from the Potato Axiom 21PT 950K screening arrays were located in CDS regions (84%), followed by 3’ UTR region (9.98%) and 5’ UTR (5.92%) (Figure S3(A)). The SIFT4G predictions categorised 152,523 sequence variants as ‘tolerated’ with a SIFT score of 1 and 24,143 variants with a score of ≤1. Annotated sequence variants comprised synonymous (43.5%), non-synonymous (39%), non-coding (15.4%) with the rest accounting for ≤ 2.1% (Figure S3(B)).

### Genotype calling and evaluation of the Potato Axiom 21PT 950K screening arrays

We performed a comprehensive evaluation of the Potato Axiom 21PT 950K screening arrays by genotyping a total of 1,247 diploid and tetraploid potato accessions. Twenty-five accessions (2.1%) were removed due to low call rate (QC call rate between 78.32% and 85.04%) and dish quality score (DQC≥ 0.82) for subsequent analyses. Successful genotype calling for 1,222 accessions led to the fitting of 834,985 sequence variants exclusive to PotatoTools Cult21PT 950K and NonCult21PT 950K clones as well as an additional 17,808 sequence variants from the Illumina Infinium 21K array (GGP3) which corresponds to 89.86% and 95.13% of the total number of sequence variants on the array (Table S5).

To ensure high-quality sequence variant selection suitable for diverse breeding and research purposes, we implemented filtering criteria on MAF (≥0.01) and SNP call rate (CR≥ 90%) on 811,515 PotatoTools Cult21PT 950K clones and Illumina Infinium 21K array (GGP3) specific variants excluding 41,278 PotatoTools NonCult21PT 950K clones specific variants (Table S5). Filtering resulted in a substantial decrease in the number of sequence variants in both PotatoTools Cult21PT 950K clones and Illumina Infinium 21K array (GGP3) specific variants. For the Illumina Infinium 21K array (GGP3), out of an initial set of 17,808 sequence variants, 14,021 variants (approximately 78.7%) passed the filtering thresholds. The PotatoTools Cult21PT 950K clones specific sequence variants, which comprised a total number of 799,210 sequence variants, a 1.37 times greater reduction to 456,639 sequence variants (approximately 57.1%) was observed compared to the Illumina Infinium 21K array (GGP3) sequence variants (Table S6).

The minor allele frequency distribution for fitted calls filtered on CR≥ 90% and MAF (≥0.01) revealed a mean MAF of 0.24 for PotatoTools Cult21PT 950K clones variants. A slightly lower value of 0.20 was observed for Illumina Infinium 21K array (GGP3) for the same criteria. Overall, sequence variants with low MAF (0.01-0.05) were more abundant than the high frequency variants with *MAF >* 0.05 in PotatoTools Cult21PT 950K clones variants (20%) suggesting a higher prevalence of rare variants (Figure S4 (A)). The proportion of the genome-wide allele missingness for the PotatoTools Cult21PT 950K clones variants was 3.12% and thus approximately 1.68 times greater than that of Illumina Infinium 21K array (GGP3) variants, which exhibited 1.85% missingness. An additional set of 22,844 sequence variants that are specific to PotatoTools NonCult21PT 950K clones filtered on CR≥ 90% were also included in the analysis described below.

### Technical reproducibility of the screening arrays

As part of the initial performance evaluation, we assessed the reproducibility of the genotyping results of the Potato Axiom 21PT 950K screening arrays by comparing sequence variant similarities between genotyping outcomes of the Agria sample that was included on each plate as a check. To assess the reproducibility of the array, 479,483 variants filtered on SNP call rate (CR≥ 90%) and MAF ≥0.01 were considered. The selected sequence variants comprised 456,639 variants specific to PotatoTools Cult21PT 950K and 22,844 from NonCult21PT 950K clones. A separate similarity analysis was performed for a total of 14,021 Illumina Infinium 21K array (GGP3) specific sequence variants. After removing missing genotype calls, an average concordance rate of 99.93% was observed for the Illumina Infinium 21K array (GGP3). A slightly lower value of 99.88% was identified for the PotatoTools Cult21PT 950K and PotatoTools NonCult21PT 950K clone specific sequence variants. For an additional quality check, we exploited the overlapping sequence variants between the SolCAP SNP array and the Potato Axiom 21PT 950K screening arrays. These 5,458 common sequence variants filtered on SNP call rate (CR≥ 90%) and minor allele frequency (MAF ≥0.01) were used for the concordance estimation of 17 potato varieties (Stich et al., 2013). On average across all 17 clones, we observed a concordance of 93% (Table S7).

### Optimising sequence variant filtering for leveraging clustering errors

We investigated all 456,639 variants specific to PotatoTools Cult21PT 950K and NonCult21PT 950K clones and an additional set of 14,021 Illumina Infinium 21K array specific variants with regards to clustering errors. The reliability of the identification of the five genotype clusters and therewith the genotype calls of sequence variants is influenced by the distance between the clusters in the two dimensions of the sequence variant cluster plot (Figure 2). To evaluate whether filtering of sequence variants can lead to an improvement of genotyping reliability, Euclidean distance and heterozygous strength offset (HetSO) were characterised. On average, the Illumina Infinium 21K array (GGP3) sequence variants showed a mean Euclidean distance of 1.10 (Figure 3A), while PotatoTools Cult21PT 950K and PotatoTools NonCult21PT 950K clones specific sequence variants exhibited a distance of 1.41 (Figure 3C). Overall, the Illumina Infinium 21K array (GGP3) sequence variants displayed narrower range of Euclidean distances (4.82e-07 to 28.16) and *σ* of 1.11, while the PotatoTools Cult21PT 950K and PotatoTools NonCult21PT 950K clones specific variants exhibited a wider range of distances, ranging from (1.94 to 29.64) with a *σ* of 1.55.

**Figure 3:**
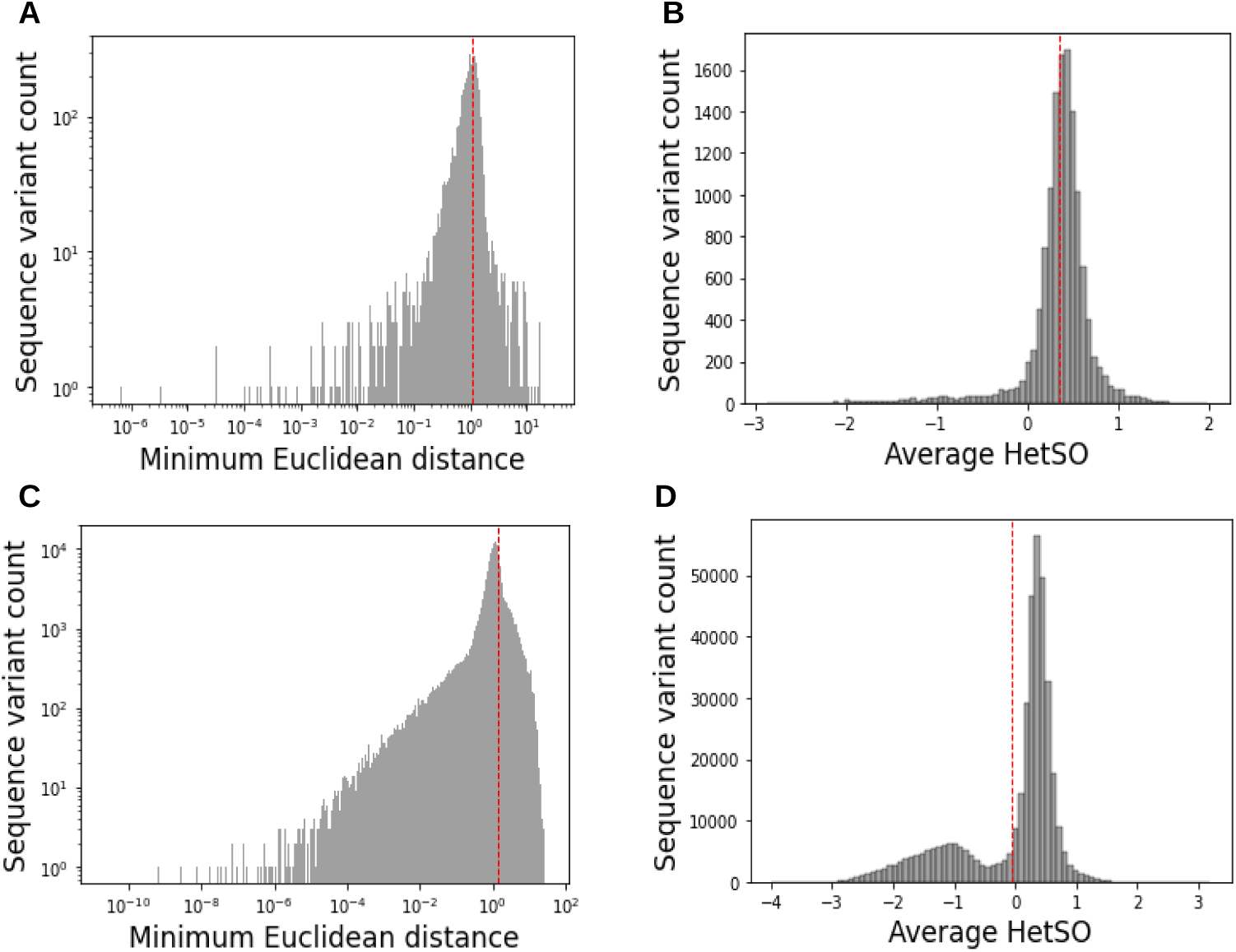
Distribution of Euclidean distance and heterozygous strength offset (HetSO) for Illumina Infinium 21K array (GGP3) specific, PotatoTools Cult21PT 950K and NonCult21PT 950K clones specific sequence variants filtered for SNP call rate (CR *≥* 90) and minor allele frequency (MAF *≥* 0.01). Histogram of (A) Euclidean distance and (B) heterozygous strength offset (HetSO) of the Illumina Infinium 21K array (GGP3), (C) Euclidean distance and (D) HetSO for the PotatoTools Cult21PT 950K and NonCult21PT 950K clones specific sequence variants. The dotted red lines represent the mean HetSO (B,D) and Euclidean distance (A,C).

An average HetSO value of 0.35 (Figure 3B) was observed for Illumina Infinium 21K array (GGP3) sequence variants and -0.05 for PotatoTools Cult21PT 950K and NonCult21PT 950K clones specific variants (Figure 3D). The HetSO values ranged from (−2.86 to 1.98) for Illumina Infinium 21K array (GGP3) specific variants with a *σ* of 0.38 and a broader range of values (−3.98 to 3.19) with a *σ* of 0.81 were exhibited by PotatoTools Cult21PT 950K and NonCult21PT 950K clones specific variants. To improve the accuracy and reliability of sequence variant calls by distinguishing between true variants and genotyping artifacts, we applied the Euclidean distance threshold of 0.05 and threshold of -0.1 for HetSO. This filtering resulted in the reduction of the number of sequence variants by approximately 1.12 times for the Illumina Infinium 21K array (GGP3) and by about 1.51 times for the PotatoTools Cult21PT 950K and NonCult21PT 950K clones specific variants (Table 1). For both Illumina Infinium 21K array (GGP3) and PotatoTools Cult21PT 950K and NonCult21PT 950K clones sequence variants, increasing the distance threshold to 0.075 and 0.1 resulted in further reductions of the number of variants. The PotatoTools Cult21PT 950K and NonCult21PT 950K clones variant, exhibited a more pronounced decrease compared to Illumina Infinium 21K array (GGP3) variants. A substantial difference in PotatoTools Cult21PT 950K and NonCult21PT 950K clones (12.38%) and Illumina Infinium 21K array (4.99%) specific variants of Cluster Type-IV indicated a significant shift in the number of variants. Conversely, PotatoTools Cult21PT 950K and NonCult21PT 950K clone variants represented a reduced number of variants in Cluster Type-V (9.36%) relative to Illumina Infinium 21K array (GGP3) variants (16.99%) (Table 1).

**Table 1:** Sequence variant categorisation into classes based on the number of clones present in the AAAA, AABB, AAAB, ABBB and BBBB clusters. Sequence variant characterisation involved 14,021 Illumina Infinium 21K array (GGP3), 456,639 PotatoTools Cult21PT 950K clones specific sequence variants filtered on (CR*≥* 90%) and (MAF *≥*0.01) and 22,844 PotatoTools NonCult21PT 950K clones specific sequence variants filtered on (CR*≥* 90%).

| Cluster class | Description | GGP3 |  |  |  | Cult21PT_950K and NonCult21PT_950K variants |  |  |  |
| --- | --- | --- | --- | --- | --- | --- | --- | --- | --- |
|  |  | Total variants | Euc=0.05, HetSo=-0.1 | Euc=0.075, HetSo=-0.1 | Euc=0.1, HetSo=-0.1 | Total variants | Euc=0.05, HetSo=-0.1 | Euc=0.075, HetSo=-0.1 | Euc=0.01, HetSo=-0.1 |
| Cluster Type-I | AAAA,BBBB,AABB,ABBB & AAAB $\neq 0$ | 7,414 | 7,175 (57.46%) | 7,136 (57.43%) | 7,105 (57.40%) | 192,803 | 182,613 (57.60%) | 181,883 (57.76%) | 181,233 (57.89%) |
| Cluster Type-II | AAAA==0 & rest $\neq 0$ | 1,386 | 1,166 (9.33%) | 1,163 (9.36%) | 1,161 (9.38%) | 43,750 | 30,335 (9.56%) | 30,197 (9.58%) | 30,101 (9.61%) |
| Cluster Type-III | BBBB==0 & rest $\neq 0$ | 1,644 | 1,398 (11.19%) | 1,394 (11.22%) | 1,391 (11.23%) | 46,206 | 35,114 (11.07%) | 35,039 (11.12%) | 34,975 (11.17%) |
| Cluster Type-IV | AAAA & BBBB==0, rest $\neq 0$ | 643 | 624 (4.99%) | 620 (4.99%) | 618 (4.99%) | 42,864 | 39,259 (12.38%) | 38,357 (12.18%) | 37,567 (12.00%) |
| Cluster Type-V | Others | 2,934 | 2,122 (16.99%) | 2,111 (16.99%) | 2,102 (16.98%) | 153,860 | 29,700 (9.36%) | 29,407 (9.33%) | 29,156 (9.31%) |
| Total |  | 14,021 | 12,485 | 12,424 | 12,377 | 479,483 | 317,021 | 314,883 | 313,032 |

### Array validation based on the proportion of incorrect calls of diploid clones

In a tetraploid genotyping space, the diploid genotypes AA, AB and BB are expected to receive the genotypes AAAA, AABB and BBBB, respectively. Deviating genotype patterns indicate errors in sequence variant genotyping. We conducted a thorough analysis of the PotatoTools Cult21PT 950K and NonCult21PT 950K clones specific sequence variants, filtered on SNP call rate (CR≥ 90%) and MAF≥ 0.01, specifically selecting 475,473 calls with at least one genotype called for diploid clones. For comparison, we also evaluated 13,981 Illumina Infinium 21K array (GGP3) sequence variants. We found 59.2% (281,655) incorrect calls for PotatoTools Cult21PT 950K and NonCult21PT 950K clones specific variants. However, also for Illumina Infinium 21K array (GGP3) sequence variants, the rate was high with 42.8% (5,987) (Table S8). The implementation of stringent filtering in SNPset-II (CR≥ 90%, MAF≥ 0.01, Hetso=-0.1 and Euclidean distance ≥ 0.05) resulted in 1.27 times fewer incorrect calls in Illumina Infinium 21K array (GGP3) (38.8%) in comparison to PotatoTools Cult21PT 950K and NonCult21PT 950K clones specific sequence variants (49.3%) (Table S8).

### Cluster shifting to correct genotype calls for diploids

Wrong genotypes of diploid clones, such as ABBB and AAAB can arise due to genotyping errors, rare variants leading to unusual allelic combinations, or the presence of interfering sequence variants close to the targeted variants. However, another possibility is that in the case of four or fewer genotype clusters, the AAAA cluster is erroneously named AAAB which in turn leads to the absence of homozygous genotypes. This situation (or the symmetrical for BBBB vs. BBBA) can be corrected by shifting the genotype clusters. The effect can be evaluated by examining if the genotype cluster shifting removed the genotype error of the diploid genotypes. We applied allelic cluster shifting to four sequence variant sets filtered based on various thresholds (Table 2), each containing varying amounts of wrong diploid genotypes. Subsequently, the allelic cluster shifting led to substantial reductions of the proportion of wrong genotype allele calls across all sequence variants. The variants filtered on MAF≥ 0.01 and CR≥ 90% exhibited the highest correction rate (74.3%). Moreover, stringent filtering on Euclidean distance and HetSO thresholds showed consistent improvements in the proportions of correct calls of 68% (Table 2) resulting in a total of 224,009 filtered sequence variants from 13 chromosomes which are designated below as high confidence sequence variants.

**Table 2:** Proportion of correct (AAAA,AABB,BBBB) and wrong allele calls (AAAB,ABBB) of diploids in a tetraploid genotyping space obtained for different filtered sequence variants sets after genotype shifting to correct genotypes for diploids. The correct and wrong calls obtained as a result of genotype shifting by applying various threshold settings i.e SNP call rate (CR), minor allele frequency (MAF), Euclidean distance (Euc distance) and heterozygous strength offset (HetSO) in four sequence variants sets.

| Description | Shifting Case-I |  | Shifting Case-II |  |
| --- | --- | --- | --- | --- |
|  | Correct calls | Wrong calls | Correct calls | Wrong calls |
| Filtered sequence variants (SNP call rate $\geq 90\%$ and MAF $\geq 0.01$) | 88,242 | 36,093 | 60,949 | 15,381 |
| Filtered sequence variants (SNP call rate $\geq 90\%$, MAF $\geq 0.01$, Euc distance $\geq 0.05$ and HetSo=-0.1) | 23,770 | 19,738 | 32,412 | 6,470 |
| Filtered sequence variants (SNP call rate $\geq 90\%$, MAF $\geq 0.01$, Euc distance $\geq 0.075$ and HetSo=-0.1) | 23,595 | 19,626 | 31,671 | 6,279 |
| Filtered sequence variants (SNP call rate $\geq 90\%$, MAF $\geq 0.01$, Euc distance $\geq 0.01$ and HetSo=-0.1) | 23,477 | 19,538 | 31,029 | 6,102 |

### Characteristics of high confidence sequence variants

The 206,616 PotatoTools Cult21PT 950K and NonCult21PT 950K clones specific high confidence sequence variants from 12 chromosomes (excluding scaffold chromosome) were evenly distributed across the Agria reference genome (dAg1 v1.0). The sequence variant density revealed an average distance of 3,629 bp between adjacent variants (median: 916 bp and *σ* of 8,131). The chromosomes were densely covered by the variants with the number of variants within 1Mb windows ranging between 0 and 890 (Figure 4A). Approximately, 27% of the sequence variants were located within 1 kilobase (kb) of a coding sequence (Table S4B). The Infinium 21K array (GGP3) sequence variants showed a lower variant density on the 12 chromosomes of the *S. tuberosum* DM v4.03 genome (Figure 4B). Across the entire genome, we observed a median inter-sequence variant distance of 3,426 bp with a *σ* of 257,531 bp.

**Figure 4:**
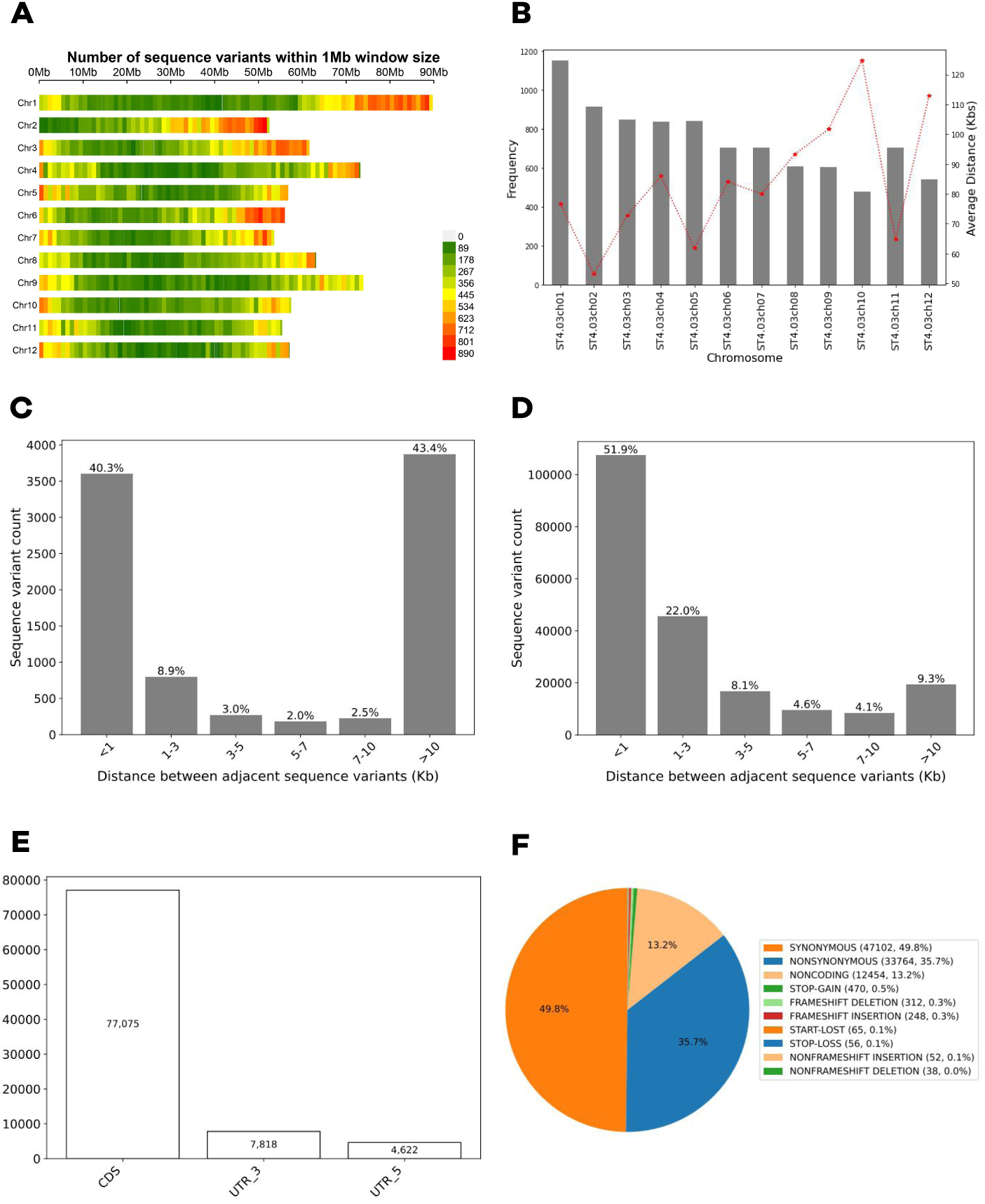
Characterisation of the filtered high quality sequence variants. (A) Heatmap of the distribution of 206,616 PotatoTools populations (NonCult21PT 950K and Cult21PT 950K clones) specific variants used for genotyping. The coloured bar represents the sequence variant density ranging from 0 to 890 per 1Mb. (B) Barplot of the number of the filtered 9,008 Illumina Infinium 21K array (GGP3) sequence variants. Dotted red lines represents the average sequence variant spacing in kilobase (Kbs) on each chromosome and grey bars represent the sequence variant count for each chromosome. (C) Frequency distribution of the distance in kilobase (kb) between adjacent 9,008 Illumina Infinium 21K array (GGP3) specific sequence variants, (D) Frequency distribution of distance in kilobase (kb) between adjacent 206,616 PotatoTools Cult21PT 950K and NonCult21PT 950K clones specific sequence variants. (E) Frequency distribution of the filtered variants of potato Axiom 21PT 950K screening array annotated in coding sequence (CDS), 3*^1^* Untranslated Region (3*^1^*UTR), and 5*^1^* Untranslated Region (5*^1^*UTR). (F) Pie-chart illustrating the functional annotation of PotatoTools Cult21PT 950K and NonCult21PT 950K clones specific sequence variants using SIFT4G.

Approximately, 51.9% of the PotatoTools Cult21PT 950K and NonCult21PT 950K specific sequence variants showed a neighbouring variation within 1kb (Figure 4D). The Infinium 21K array (GGP3) sequence variants showed 40.3% of variants within 1kb (Figure 4C). We were able to predict amino acid substitutions for a total of 94,561 PotatoTools Cult21PT 950K and NonCult21PT 950K clones specific variants. The genomic position-based annotation revealed that 84.5% of the variants were located in the CDS (Figure 4E) of which 75.3% were tolerated mutations and 9.22% deleterious. A total of 13.5% of the variants were located in the UTR regions. Moreover, 49.8% of the variants were labelled synonymous, 35.7% non-synonymous, 13.2% non-coding, the rest being other variants (Figure 4F).

### Application of Potato Axiom 21PT 950K array for population structure, genome-wide association, and genomic prediction analysis

A total of 215,496 sequence variants comprising 9,008 Illumina Infinium 21K array (GGP3) and 206,488 PotatoTools Cult21PT 950K clones specific variants from the Potato Axiom 21PT 950K arrays were used to characterise the population structure in a set of 1,222 unique potato clones. In the first step, the clones of the resequencing panel were considered in the PCA analysis of diploids, non-tuberosum diploids and tetraploid clones. The first principal component (PC1) accounted for 4.51% of the total variance and separated the two ploidy levels (diploids and tetraploids) of potato, whereas the second principal component (PC2) accounted for 2.66% of the total variance (Figure 5A). Moreover, a subcluster consisting of the clones H89 5771 and P40 (Figure 5A) was revealed in the analysis. In a second step, we evaluated the population structure of the tetraploid clones of the resequencing panel based on the country of origin. In this case, we observed an intermixing of the clones associated with different countries of origin (Figure 5B). In this particular case, PC1 and PC2 accounted for 2.92% and 2.61% variance, respectively. Furthermore, we also assessed the population structure of the tetraploid clones from the resequencing panel and PotatoTools populations but excluded 30 genotypes that are genotyped multiple times. The first two principal components accounted for 1.5% and 1.4% of the total sequence variant variability. Overall, the clones in PCA analysis clustered together exhibiting clear population structure of clones according to their market segment with a few exceptions (Figure 6). The PCA analysis did not reveal any separation between PotatoTools populations and resequencing panel-specific clones.

**Figure 5:**
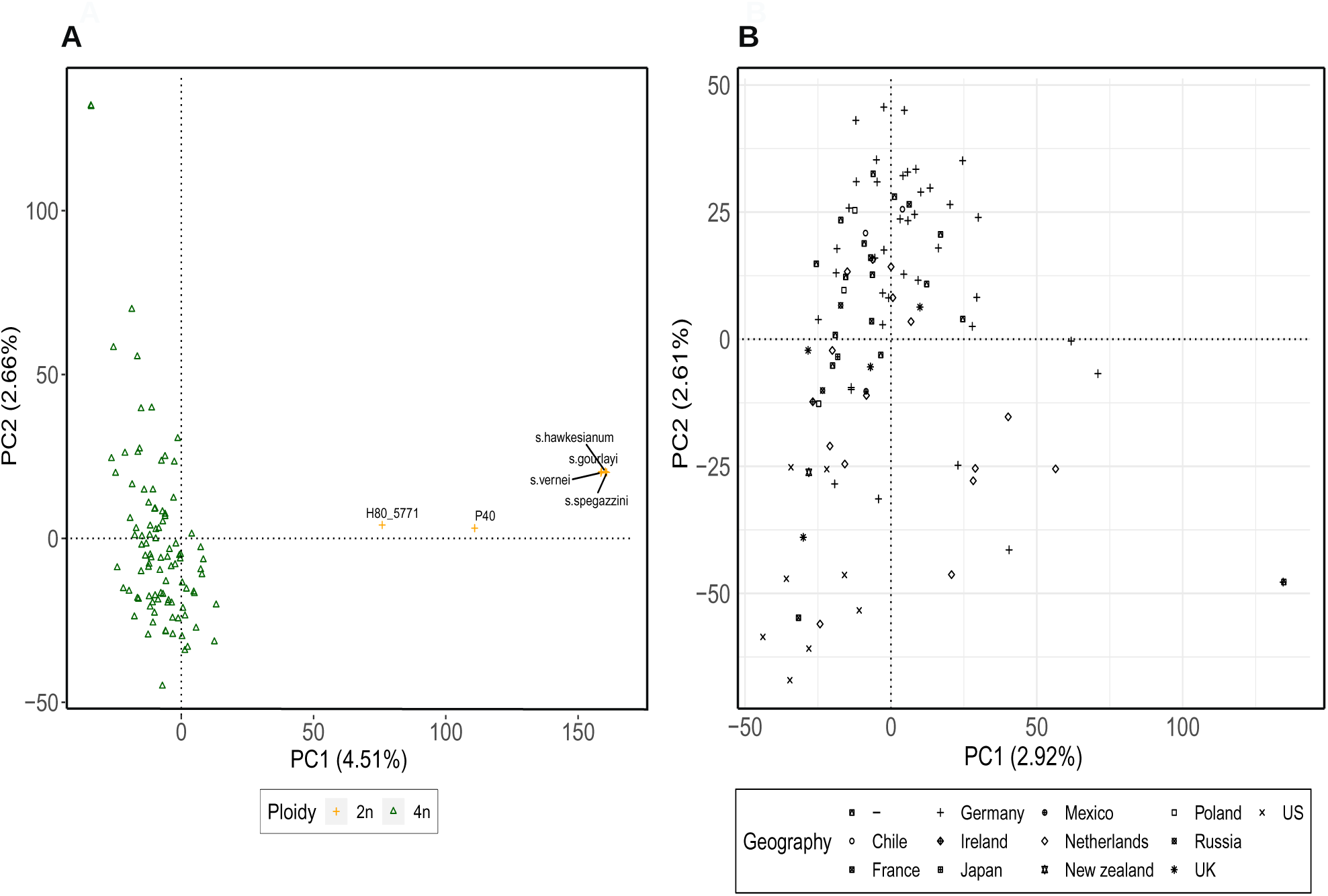
Principal component analysis (PCA) of all resequencing panel based on 215,496 sequence variants. (A) PCA of resequencing panel clones, and (B) PCA of the tetraploid clones from resequencing panel, where PC1 and PC2 refers to the first and second principal components, respectively. Numbers in parenthesis refer to the proportion of total variance explained by each principal component.

**Figure 6:**
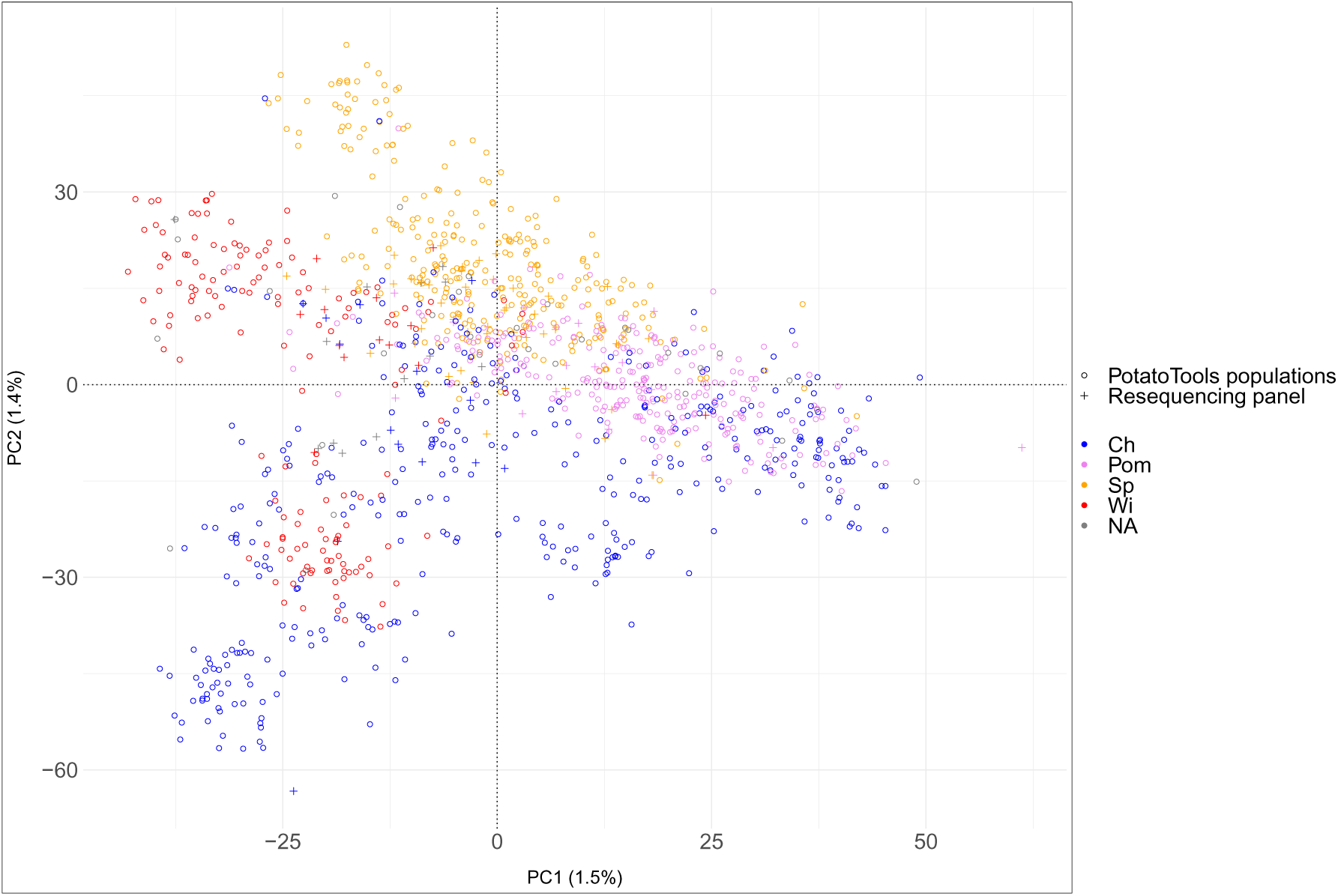
Principal component analysis (PCA) of the PotatoTools populations and resequencing panel. Principal component analysis of 1,192 clones belonging to a diverse set of market segments using 215,496 sequence variants. PC1 and PC2 refer to the first and second principal components, respectively. Numbers in parenthesis refers to the proportion of total variance explained by each principal component. Each clone is represented by a different colour corresponding to the market segments. Four market segments namely chips/crisp (Ch), french fries (Pom), starch (Wi), and table (Sp) were used for the grouping of the entries in the PCA. Genotypes with missing information about market segmentation were represented with NA. The shapes of the dots represent the genotype origin being the resequencing panel or the PotatoTools populations.

In the DAPC analysis, the minimum Bayesian information criterion (BIC) of the k-means clustering algorithm suggested five genetic groups. The DAPC showed genetic sub-structure corresponding partly to market segmentation categories by the first four discriminant functions (Figure 7). The first discriminant function (DF1) explained 37.8% of the variance, while the second function (DF2) accounted for 27.6%. Two clusters, corresponding to table processing (Sp) and starch (Wi) clones, were separated in the discriminant space. Within the table processing clones, a distinct sub-cluster was observed that overlapped with genotypes from other market categories. The remaining clusters showed extensive overlap and contained a mixture of clones from all market segmentation categories, indicating genetic affinity among the clones.

**Figure 7:**
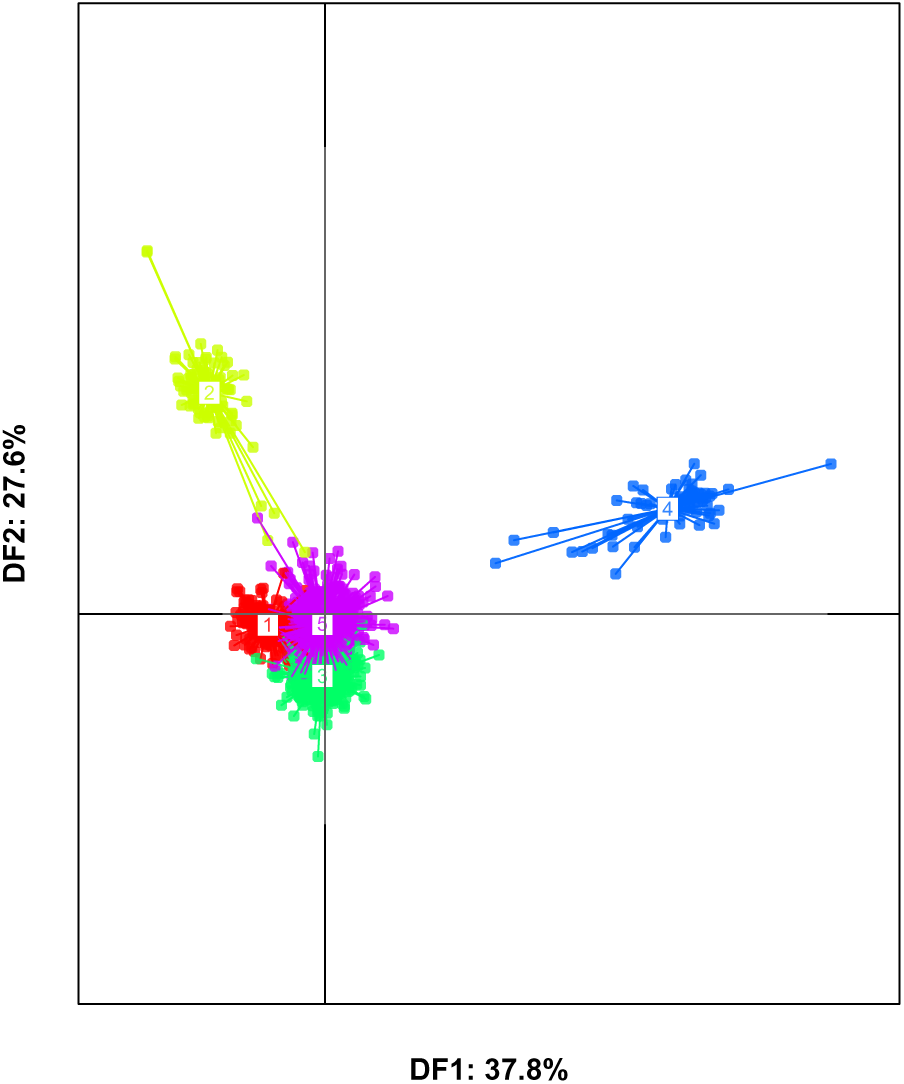
Discriminant analysis of Principal Components (DAPC) of 1,192 PotatoTools and resequencing panel clones based on 215,496 sequence variants. Scatterplot of the first two discriminant functions (DF1) and (DF2), which explain 37.8% and 27.6% of the total variance, respectively, Each point represents an individual genotype, and colours indicate the assigned market segmentation cluster identified by DAPC. The cluster 1 represents chips/crisps (Ch), 2 represents starch (Wi), 3 represents french fries (Pom), 4 and 5 represents table variety (Sp).

GWAS was performed to explore the genetic basis of polyphenol oxidase (PPO) activity in potato. The phenotypic data of 998 potato clones as well as 194,621 markers filtered on missing values higher than 20% and those with MAF below 0.05 were used for the analysis. The sequence variants from the Chromosome Unknown of PotatoTools Cult21PT 950K and NonCult21PT 950K, Chr00 from Illumina Infinium 21K array (GGP3) and Chloroplast were removed from the analysis due to their uncertain genomic positions and distinct evolutionary origin with lower variability, respectively. Furthermore, we aligned the flanking sequence of sequence variants from Illumina Infinium 21K array (GGP3) on Agria reference genome (dAg1 v1.0) using the Basic Local Alignment Search Tool (BLAST) to obtain their positions on this reference genome.

We identified 84 PotatoTools Cult21PT 950K and NonCult21PT 950K clones specific significant sequence variants associated to PPO with − log_10_ p-values *>* 6.59 with a notable peak observed between 51-54 Mb on chromosome 8 (Figure 8A and 8B). These sequence variants were annotated to 35 unique genes (Figure 8C) and (Table S10). Moreover, 4 significant Illumina Infinium 21K array (GGP3) were also present in the peak region annotated to 2 unique genes highlighted in red (Figure 8C) and (Table S11).

**Figure 8:**
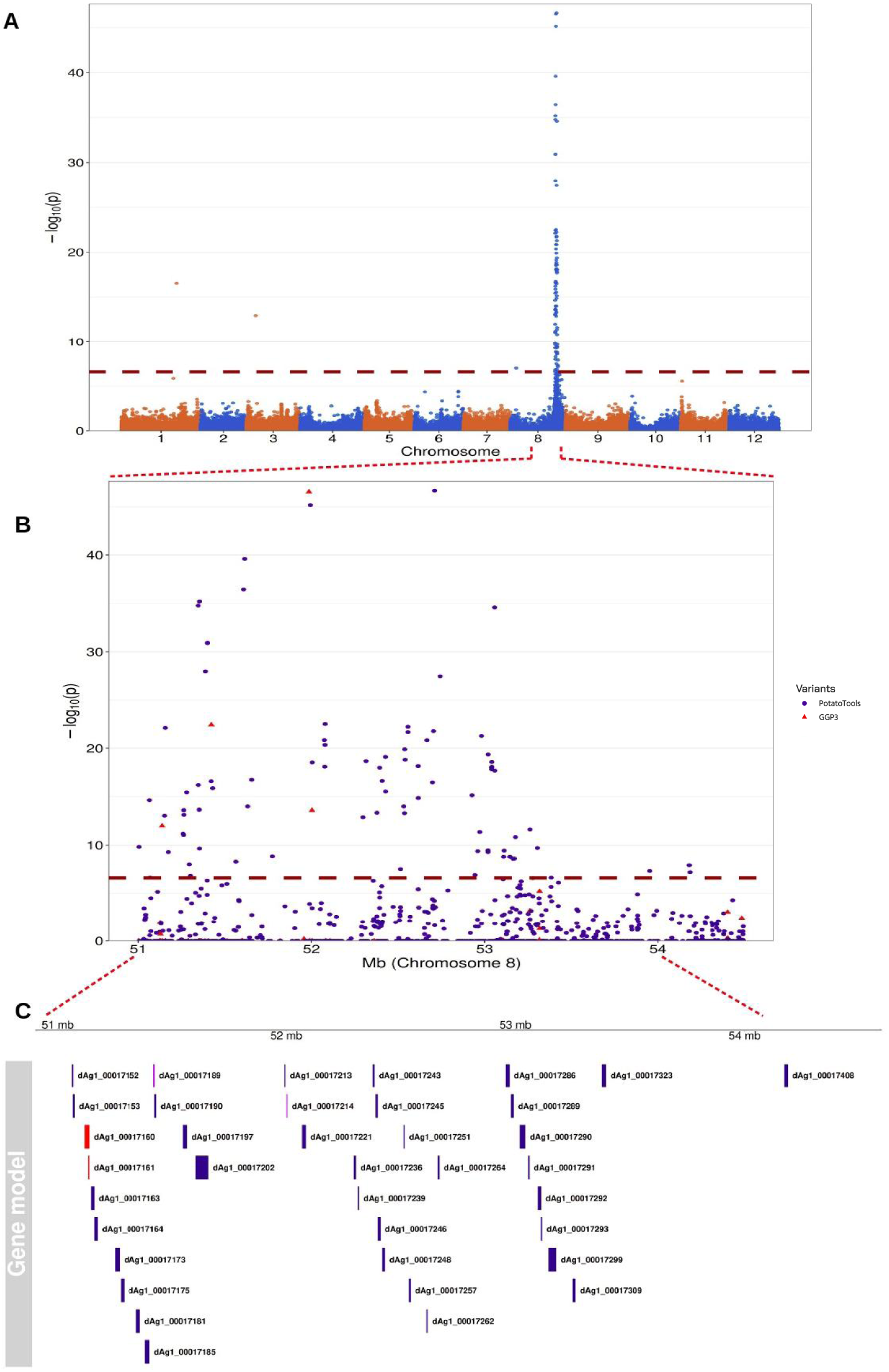
Genome-wide Manhattan plot for the genome-wide association analysis of sequence variants of Axiom 21PT 950K screening array for 988 clones for polyphenol oxidase (PPO) activity. (A) The x-axis represents the chromosomes of the diploid Agria genome (dAg1 v1.0) and y-axis corresponds to the negative logarithm of the p-values (*−* log_10_ p-values). The horizontal dashed line indicates the genome-wide significance threshold. (B) Zoomed-in view for the 51-54 Mb region on Chr08, showing a finer resolution of the peak region. (C) Gene models within the 51-54 Mb region corresponding to the GWAS peak, highlighting candidate genes depending on the SNP set causing significance (red=Illumina Infinium 21K array (GGP3), blue=PotatoTools NonCult21PT 950K, and Cult21PT 950K clones and purple=both).

Prediction accuracy, measured as the correlation between observed and predicted phenotypic values, ranged from 0.71 to 0.86 (mean 0.79) for the Illumina Infinium 21K array (GGP3) specific variants indicating a high reliability. In contrast, the PotatoTools Cult21PT 950K and NonCult21PT 950K clones specific sequence variants achieved predictive abilities ranging from 0.72 to 0.86 with a slightly higher mean accuracy of 0.80.

## DISCUSSION

Sequence variants such as SNPs are nowadays important tools in plant as well as animal breeding programmes (Ganal et al., 2012, Rasheed et al., 2017). Depending on the aim, two strategies can be distinguished: assessing the allele configuration (i) at selected loci or (ii) across the entire genome. Different genotyping techniques are dominantly used for each of these strategies. For genotyping individual or selected sequence variants e.g. marker-assisted selection programmes of mono- or oligogenic traits, the Kompetitive Allele Specific PCR (KASP) (Semagn et al., 2014) is to our knowledge the technique of choice used in commercial plant breeding programs. The alternatives applied for genome-wide characterisation of allele configuration are skim sequencing (Kumar et al., 2021), genotyping by sequencing (GBS) (He et al., 2014), Potato Multi-allele scanning haplotags (PotatoMASH) (Leyva-Pérez et al., 2022), and methods such as targeted genotyping-by-sequencing or exploitation of hybridisation-based genotyping methods such as SNP arrays (You et al., 2018, Rasheed et al., 2017). In the context of commercial breeding programmes, the use of SNP arrays is preferred due to their easier integration in high throughput pipelines (You et al., 2018), the reasons for this being they do not need computation intensive bioinformatic analyses such as read mapping, variant calling and filtering pipelines. Moreover, SNP array-based data analyses frequently do not require imputation of missing data (Hussain et al., 2017).

In contrast, the development of efficient SNP genotyping arrays is a significant challenge, especially for autopolyploid genomes such as that of potato. The existing potato arrays SolCAP Array and Illumina Infinium Potato Array (Hamilton et al., 2011, Felcher et al., 2012, Vos et al., 2015, Hardigan et al., 2017) have been widely used in breeding and research including population genetic, but also quantitative genetic analyses (Yousaf et al., 2023, Jo et al., 2022, Pandey et al., 2021, Stich et al., 2013, Ellis et al., 2018, Alvarez-Morezuelas et al., 2023, Klaassen et al., 2019). However, the usage of limited genotypes to identify the sequence variants and low variant density are the major drawbacks of these arrays. Therefore, our study focused on the development of a potato array based on advanced read technology, improved reference genome, more diverse set of genotypes with better sequence variant coverage.

In this study, we developed two high-density Axiom-based SNP genotyping screening arrays for potato, using 10X based linked reads technology and the dAg1 v1.0 reference genome (Freire et al., 2021). The primary goal behind the design of this array was to provide a robust and high-density genotyping solution for applications in commercial potato breeding programmes but also for publicly funded research projects. To date, Potato Axiom 21PT 950K screening arrays are the first reported high-throughput SNP genotyping arrays incorporating 947,845 sequence variants identified using a sequence variant discovery panel of 108 diverse potato clones representing worldwide diversity, market segment, and disease resistance traits. Additionally, the incorporation of sequence variants from the Illumina Infinium array and SolCAP array allows the integration of previously generated data. Furthermore, our arrays provide the following advantages: First, the arrays were developed using the largest resequencing data set available at that time, providing increased statistical power for variant detection and ensuring a comprehensive representation of the genetic diversity, thereby giving robust and accurate genetic association studies. Secondly, our arrays included potato clones from diverse geographical regions compared to previous arrays where in the case of SolCAP, sequence variants were identified to a major extent from the North American varieties bred for use in the processing industry (Hamilton et al., 2011). Available arrays struggle with capturing rare variants, therefore hindering identification of allelic segments from distantly related genetic resources and causing ascertainment bias (Moragues et al., 2010). Lastly, the three-tier strategy, starting from sequence variant identification, followed by filtering, probe development based on whole genome sequencing accessions and final array evaluation reduces the risks of selection of false positive variants in our arrays which is discussed below.

### Array design and genotyping assessment

Across 98% of the samples, 90% of the sequence variants unique to PotatoTools resequencing panel (NonCult21PT 950K and Cult21PT 950K clones) were successfully genotyped suggesting a high genotyping power. Overall, the proportion of successfully genotyped variants were higher in our study compared to the values previously reported i.e 80.8% in Vos et al. (2015), 45% in Hirsch et al. (2013) and 38% in Lindqvist-Kreuze et al. (2014) for potato. The higher percentage of successfully genotyped sequence variants in our arrays is a result of their careful selection in our study. This explanation is in agreement with the observation that for the sequence variants originating from the Illumina Infinium 21K array (GGP3), which underwent one additional round of stringent filtering and selection for better performance an only slightly increased genotyping rate of 95% was observed.

The technical array reproducibility of our Potato Axiom 21PT 950K screening arrays showed less than 1% variation which is in line with previously reported values for arrays of potato (Vos et al., 2015), *Cicer arietinum* (Roorkiwal et al., 2018) and *Glycine max* (Lee et al., 2015). Issues such as sample contamination, partial probe binding to secondary homologous sites, or errors in genotype calling software when identifying the correct genotype clusters might be the reason for this remaining variability. Moreover, laboratory or sampling issues might also contribute to some level of variability. Low-quality DNA was the reason for high levels of discordance in the *Axiom J. regia* 700K array (Marrano et al., 2019). Furthermore, genotyping using fitPoly appeared to be significantly more accurate than GenomeStudio, where a 1.7% discrepancy has been reported for genotype calls for both diploid and tetraploid clones (Hirsch et al., 2013).

Our results depict a high degree of SNP reproducibility (93%) between the genotyping profiles of a common set of 17 clones based on the SolCAP SNP array and our Potato Axiom 21PT 950K screening arrays suggesting that our genotyping procedure works well. The observed variability of 6.9% is presumably caused by usage of the same genotype of different origins and DNA quality which can lead to discrepancies in reproducibility.

The stringent filtering applied in our study based on SNP call rate and minor allele frequency resulted in 493,504 high-quality sequence variant calls for each clone. This filtering was used as a way to remove false positives, thereby improving the performance of the selected variants in applications such as population genetic studies, genetic mapping, and genomic prediction. The SNPolisher’s (v.1.5.2) “Ps Classification” function that is frequently used in studies with SNP genotyping in diploid species (Howe et al., 2020, Montanari et al., 2019) and allopolyploids (Bassil et al., 2015) for classifying sequence variants into cluster categories according to their reliability e.g Cluster Type-I to Cluster Type-V (Figure 2) does not apply to autopolyploids. Therefore, we designed a custom scheme for stringent variant filtering by inspecting the Euclidean distance among the clusters and HetSO in two dimensions (Figure 2) to reduce the genotype clustering errors. With this procedure, we selected 329,506 probes where allele dosage can be inferred reliably.

Our study design allowed us to filter the sequence variants based on the genotypes of the available diploid samples that are unexpected such as AAAB or ABBB and are considered as potential indicators of genotyping errors (Vos et al., 2015). We detected 59% of the sequence variants from the PotatoTools NonCult21PT 950K and Cult21PT 950K specific variant (Table S8) calls for diploids that are not possible in a tetraploid genotyping space. This percentage was higher than the 43% observed for the Illumina Infinium 21K array (GGP3) specific sequence variants. The addition of diploid genotypes proved to help identify and remove poor sequence variant calls, whereas, in the study of Vos et al. (2015), genotype calling without the diploid samples was used for further analysis as the inclusion of diploids resulted in too many rejected markers. However, because of the large number of available sequence variants, we decided for the conservative approach and removed those variants from further analysis. As indicated above, the lower percentage of wrong calls in the Illumina Infinium 21K array (GGP3) can be explained by the pre-selection of markers on previous potato arrays.

Instead of removing all wrongly called genotypes or completely removing diploids from the analysis, we carefully examined and re-assigned allele calls by applying cluster shifting. It was applied to those cases where missing genotype clusters led to mis-genotyping of diploids. By systematically shifting the genotype calls, we were able to rescue 56,182 variants, resulting in a total of 224,009 high-quality filtered sequence variants suitable for the design of the Potato Axiom production array.

In our genomic analysis, we found that the filtered sequence variants tended to be close to genes, with notable proportions (67.7%) within 1-4 kilobase (kb) (Table S4B). On average, the sequence variants maintained an overall average distance of approximately 3,629 bp and a median of 916 bp. With these numbers in the context of an LD decay with 265 bp to *r*^2^ or *D^t^* of 0.1 (Stich et al., 2013), we are confident that our array will be a powerful tool for GWAS and genomic prediction. To illustrate this, we assessed the applications of the array for such analyses.

### Applications of the Potato Axiom 21PT 950K array

The PCA analysis of the resequencing panel demonstrated the clear separation of diploid and tetraploid clones along the first principal component (Figure 5A). This is in accordance with the results of the study of Stich et al. (2013). In the present study, no distinct clusters were visible in the PCA analyses based on sequence variants of tetraploid clones belonging to diverse geographical regions (Figure 5B). This result is analogous to the findings of previous studies (Selga et al., 2022) of the Nordic potato breeding programme. No evident clusters might be the result of the recombination of genetic material of diverse origins in the context of commercial breeding programmes. However, the history of the potato domestication and selective breeding for traits such as yield, taste, and disease resistance also has the potential to shape diversity among potato genotypes. This is in agreement with the observation of a separation of clones belonging to different market segments. Our DAPC analysis revealed a clear separation between certain market segmentation groups (e.g., table processing vs. starch varieties) and substantial overlap among others, indicating weak but structured diversity (Figure 7). Similar clustering patterns were reported by Deperi et al. (2018),D’hoop et al. (2010), and Wilson et al. (2021), where market segmentation-based clustering showed limited genetic separation. The market segmentation categories do not always align with the underlying genetic structure due to extensive gene flow, shared breeding lines across market classes, and selection for similar agronomic traits across market groups.

In this study, we demonstrated the potential of the Potato Axiom 21PT 950K testing arrays for GWAS and furthermore for genomic prediction of PPO activity in potato. Tuber bruising resulting from mechanical impact on tubers triggers phenolic oxidation by polyphenol oxidases (PPO) which in turn produces melanin pigments which are responsible for tuber browning (Thygesen et al., 1995, Urbany et al., 2011, Wright et al., 2005). This trait is vital for determining the visual quality of potatoes, particularly in table potatoes exhibiting higher starch content. Moreover, in processed products like fries and crisps, where reduced browning is required.

The classical tuber bruising score requires a destructive analysis of several kilos of tubers. Therefore, the assessments of PPO activity have been suggested for breeding programmes as indirect selection characters which are correlated with tuber bruising susceptibility (Urbany et al., 2011). Previous research identified several candidate genes and Quantitative Trait Loci (QTLs) linked to bruising on chromosome 8 (Angelin-Bonnet et al., 2023, Urbany et al., 2011). Notably, the POT32 gene, a key gene present on chromosome 8, has been identified along with other genes influencing PPO activity in tubers (Urbany et al., 2011).

The analysis of the significant variants identified in Illumina Infinium 21K array (GGP3) and PotatoTools NonCult21PT 950K and Cult21PT 950K clones specific variants on chromosome 8 provided a deep understanding of their contributions and biological implications in PPO activity (Table S10 and S11). Having a closer look into the identified peak region between 51–54 Mb, we were able to find 85 significant variants in 37 genes.

A BLAST analysis was carried out to further validate the functional relevance of the identified genes. In particular, SOLTUB.AGRIA.G00000028441 showed 100% similarity to *Solanum tuberosum* tuber polyphenol oxidase PPO (POT32 allele) gene and SOLTUB.AGRIA.G00000028442 to *Solanum tuberosum* tuber polyphenol oxidase PPO (POT33 allele) gene. These results aligned with previous findings of Werij et al. (2007), where molecular markers from POT32, have been found to co-localize with a QTL associated with the enzymatic discolouration on potato chromosome 8 in a diploid mapping population. This finding illustrated that the identified associations are biologically meaningful, supporting the relevance of the associations identified on chromosome 1 at 62,543,122 bps (dAgria v1.0 Chr01 176,032) and dAgria v1.0 Chr03 546,095 at 10,578,483 bps on chromosome 3.

It is noteworthy that, using genomic prediction, we were able to capture and explain more genetic variance compared to GWAS variants, as genomic prediction uses genomewide sequence variants information rather than focusing on a limited set of significant markers.

Our Axiom 21PT 950K array offers several key advantages over the Illumina Infinium 21K array (GGP3) specific variants, making it a powerful tool for both GWAS and genomic prediction. Overall, it provides higher marker density and improved uniform genome-wide coverage. These improvements are critical for the detection of causal loci and fine-mapping resolution in GWAS while also increasing the prediction accuracy of genomic prediction. Looking ahead, the implications of the Axiom 21PT 950K array are far-reaching. Our potato array will accelerate the potato breeding programmes by enabling the selection of variants linked with key traits, deriving the development of better potato varieties with optimised traits such as yield, disease resistance and tuber quality.

## ACKNOWLEDGEMENTS

Computational infrastructure and support were provided by the Center for Information and Media Technology at (ZIM) at Heinrich Heine University Düsseldorf.

## AUTHORS CONTRIBUTION STATEMENTS

Nadia Baig was involved in conceptualisation, data analysis and writing. Kathrin Thelen and Mathieu A.T.Ayenan carried out the GWAS and genomic prediction analysis. Vanessa Prigge, Stefanie Hartje, Katja Muders, Juliane Renner, Bernd Truberg, Julien Bruckmüller, Jens Lübeck, Evelyn Obeng-Hinneh, Rafal Zgadzaj, Arne Rosen contributed to funding acquisition, genetic and phenotypic material collection, and review. Delphine van Inghelandt was involved in conceptualisation, funding acquisition, and review. Benjamin Stich was involved in conceptualisation, funding acquisition, project design, project coordination, writing and review.

## DATA AVAILABILITY

The data used in the current study are part of the corporate company SaKa Pflanzenzucht GmbH & Co. KG, Nordring-Kartoffelzucht und Vermehrungs-GmbH & Co. KG and EUROPLANT Innovation GmbH & Co. KG proprietary database and cannot be shared due to privacy and confidentiality restrictions. However, encoded versions of the data can be provided upon request. The analysis codes are also available upon request.

## ETHICS APPROVAL AND CONSENT TO PARTICIPATE

The authors declare that the experimental research done on the plants mentioned in this paper complied with institutional and national guidelines.

