## Supplementary Data for "An Axiom SNP genotyping array for potato: development, evaluation and applications"

### SUPPLEMENTAL MATERIALS

#### Method S1: Sequence variant annotation

The SIFT4G database was built using genomic DNA sequence in FASTA format, the gene annotation file (GTF), and protein sequence file. The amino acid substitutions having SIFT median score $\leq$0.05 were considered deleterious, while variants with score $>$0.05 were categorised as tolerated.

#### Method S2: Initial sequence variant selection

The overall goal of this step was to select 32,000 sequence variants from PotatoTools NonCult21PT_950K, and 2.85 million from PotatoTools Cult21PT_950K clones for which the Affymetrix *p_convert* value is calculated.

The sequence variants were selected based on the following procedure which was designed to ensure a uniform distribution of variants across the genome, but at the same time an increased number of variants in gene dense regions:

1) Initially, the potato reference genome was divided into 25kb windows and further supplemental 5kb segments per window for the PotatoTools Cult21PT_950K clones specific variants.

2) The number of sequence variants required in each of the two window types was calculated:

i) 25Kb bins: Gene density ratio was calculated by the following equation (1):

$$\mathrm{GD}_{ratio} = \frac{\frac{1}{2}S_T}{G_T}, \tag{1}$$

where, $S_T$ was the number of sequence variants to be selected for scoring and $G_T$ was the total number of genes in the annotation file. Later on, $GD_{ratio}$ was multiplied with the number of genes present in each 25 kb bin to identify the number of required sequence variants.

ii) For 5kb supplemental bins, the number of required sequence variants was calculated by equation (2):

$$R_{snps} = \frac{\frac{1}{2}S_T}{T_{bins}} \tag{2}$$

where, $S_T$ was the total number of sequence variants to be selected for scoring and $T_{bins}$ was the total number of 5kb bins. According to the Thermofisher Scientific's SNP array design instructions, sequence variants having a higher number of interfering mutations (wobble count) are less likely to be successfully converted to an informative sequence variant and, thus obtain a high *p_convert* score. Therefore, we a applied sequence variant prioritisation scheme based on the number of interfering sequence variants to the target sequence variants.

3) Priority assignment was performed for all sequence variants by considering the distance of the target sequence variant to the neighbouring sequence variant. The sequence variants having no interfering marker at a distance of 35 bp's were grouped into (category I) and assigned a highest priority P1. Those with interferring marker at 34 bp distance as second higest priority P2. Priorities were subsequently reduced in accordance with decreasing distances until reaching P35, where the distance was reduced to 1 bp between the target and its neighbouring sequence variant (category II). Furthremore, the ones exhibiting two interfering markers were given priorities from P36 to P103. Similar criteria of prioritisation were followed for the target variants having more than 2 neighbouring sequence variants (category IV). This selection criteria were applied to ensure that the selected variants provide independent information about the underlying genetic variation in the population.

4) To increase the likelihood of selecting informative sequence variants, the prioritisation scheme

also considered minor allele frequency (MAF), sequence variant class, distance based priority value, and type of sequence variant alleles. A Python based script was developed to assign weights to the following selection criteria:

(i) The sequence variants prioritised as P1 to P35 from step (3) were given weights that decreased incrementally, with P35 assigned a weight of 1000 with each preceding sequence variant of less priority receiving a slightly lower weight. (ii) Sequence variants were further classified into minor allele frequency (MAF) classes. A weight of 25 was assigned to sequence variants having $0.05 \leq$ MAF$< 0.15$, 50 to the sequence variants with $0.15 \leq$ MAF$< 0.25$, and a weight of 75 for sequence variants with $0.25 \leq$ MAF$< 0.35$ , 100 and 125 was assigned to variants having $0.25 \leq$ MAF$< 0.35$, and MAF$\geq 0.45$ respectively.

(iii) All A/T and C/G sequence variants and indels were assigned a weight of 1. Sequence variants other than A/T, C/G alleles and biallelic sequence variants were given a weight of 100. All weights were summed for respective attributes for the sequence variants. The number of required sequence variants in equation (1) and equation (2) was calculated and selected.

5) For PotatoTools NonCult21PT_950K clones specific variants 150kb bins were made. In bins with both intergenic and genic mutations, one sequence variant was selected from each category without replacement. For bins exclusively containing intergenic mutations, only one variant was selected. The selected sequence variants contained private alleles specific to each uncultivated clone. The final list of selected sequence variants from the above described steps along with the 21,227 markers from the Illumina Infinium 21K array (GGP3) array were submitted to Affymetrix for evaluation if the sequence variants can be tiled, which is realized by calculating the *p_convert* value. The *p_convert* value takes a value between 0 and 1 to describe the predicted probability to tile the sequence variant on array using its sequence, binding energies, degree of non-specific binding

and hybridisation to multiple genomic regions *p_convert* score categorised sequence variants as "recommended", "neutral", "not recommended", and "not possible". Sequence variants tagged as "recommended","not recommended" and, "neutral" were further analysed for array tiling.

**Table S1: Accessions of the resequencing panel used for sequence variant identification and their year of release, country of origin, ploidy, and market usage.**

| Name | Year of release | Country | ploidy | Market usage | Name | Year of release | Country | ploidy | Market usage |
|---|---|---|---|---|---|---|---|---|---|
| Adretta | 1975 | Germany | 4n | Puree and soups | Lilly | 2011 | Germany | 4n | Salad making |
| Agria | - | Germany | 4n | Multi-purpose | Marabel | 2001 | Germany | 4n | Fries processing |
| Albatros | 1996 | Germany | 4n | Starch processing | Maris Piper | 1967 | UK | 4n | Chip processing |
| Allians | 2003 | Germany | 4n | Salad making | Natalia | - | - | 4n | Fries processing |
| Altus | 2008 | Netherlands | 4n | Salad making | Nevsky | - | Russia | 4n | Salad making |
| Ambition | 2008 | Netherlands | 4n | Multi-purpose | Nicola | 1973 | Germany | 4n | Salad making |
| Atlantic | 1976 | US | 4n | Chip processing | Odysseus | 2018 | Germany | 4n | Crisps processing |
| Atzimba | - | Mexico | 4n | Fries processing | Olympus | - | UK | 4n | Crisps processing |
| Belana | 2000 | Germany | 4n | Salad making | Ona | - | Chile | 4n | Salad making |
| Bintje | 1910 | Netherlands | 4n | Fresh market | Otolia | 2014 | Germany | 4n | Fresh market |
| BNA1 | - | Germany | 4n | Starch processing | P3 | - | - | 2n | - |
| BNA2 | - | Germany | 4n | Fresh market | P40 | - | - | 2n | - |
| BNA3 | - | Germany | 4n | - | Pentland Dell | - | UK | 4n | Fries processing |
| BNA4 | - | Germany | 4n | Fresh market | Pirol | 2000 | Germany | 4n | Crisp processing |
| BNA5 | - | Germany | 4n | Starch processing | Premier Russet | - | US | 4n | Fries processing |
| Cara | 1973 | Ireland | 4n | Multi-purpose | Princess | 1997 | Germany | 4n | Salad making |
| Celtiane | 2008 | France | 4n | Fresh market | Quadriga | 2005 | Germany | 4n | Starch processing |
| CGN17888-16 | - | - | 2n | - | Quarta | - | Germany | 4n | Multi-purpose |
| CGN18114-14 | - | - | 2n | - | Record | 1966 | UK | 4n | Crisps processing |
| Charlotte | 1981 | France | 4n | Fries processing | Regina | 2009 | Germany | 4n | Salad making |
| Cherie | 1997 | France | 4n | Multi-purpose | Rode Eersteling | - | Netherlands | 4n | Salad making |
| Colomba | - | Netherlands | 4n | Multi-purpose | Rooster | - | Ireland | 4n | Multi-purpose |
| Dark Red Norland | 1999 | US | 4n | Multi-purpose | Rudolph | 2008 | Netherlands | 4n | Multi-purpose |
| Desiree | 1962 | Netherlands | 4n | Multi-purpose | Russet Burbank | 1917 | US | 4n | Multi-purpose |
| Donata | 2013 | Germany | 4n | Fries processing | SA222-2 | - | - | 2n | - |
| Early Rose | - | - | 4n | Multi-purpose | SA225-2 | - | - | 2n | - |
| Edison | 2017 | Germany | 4n | Fries processing | Saskia | - | Germany | 4n | Multi-purpose |
| Eurogrande | 2009 | Germany | 4n | Starch processing | Schwalbe | 1956 | Germany | 4n | Multi-purpose |
| Europrima | 2006 | Germany | 4n | Starch processing | Semlo | - | Germany | 4n | Starch processing |
| Felsina | - | Netherlands | 4n | Fries processing | Seresta | 1995 | Netherlands | 4n | Multi-purpose |
| Flava | - | Germany | 4n | Crisp processing | SH C909 | 2018 | Netherlands | 4n | Fries processing |
| Fontane | 1999 | Netherlands | 4n | Fries processing | Shepody | 1980 | US | 4n | Fries processing |
| Gala | 2002 | Germany | 4n | Multi-purpose | Skawa | 2000 | PL | 4n | Starch processing |
| Gladiator | 1994 | New zealand | 4n | Fries processing | Snowden | 1990 | US | 4n | Chip processing |
| GLKS30976-20 | - | - | 2n | - | Solist | 1999 | Germany | 4n | Fresh market |
| Granola | 1975 | Germany | 4n | Multi-purpose | Spunta | 1967 | Netherlands | 4n | Multi-purpose |
| H98.D12 | - | - | 4n | - | Talent | 2006 | Germany | 4n | Fries processing |
| Harpun | 1993 | Poland | 4n | Multi-purpose | TOYOSHIRO | 1976 | Japan | 4n | Crisp processing |
| Hermes | 1973 | Netherlands | 4n | Multi-purpose | Udacha | - | Russia | 4n | Multi-purpose |
| Innovator | 1999 | Netherlands | 4n | Fries processing | Velox | 1994 | Germany | 4n | Multi-purpose |
| Jelly | 2002 | Germany | 4n | Fries processing | Verdi | 2003 | Germany | 4n | Crisp processing |
| Jukijiro | 1961 | Japan | 4n | Salad making | Vitabella | 2011 | Netherlands | 4n | Multi-purpose |
| Karelia | 2016 | Germany | 4n | Starch processing | VR808 | - | Netherlands | 4n | Crisp processing |
| Karolin | 1990 | Germany | 4n | Multi-purpose | Yagana | - | Chile | 4n | - |
| Kathadin | 1932 | US | 4n | Multi-purpose | Zorba | 1997 | Germany | 4n | Fries processing |
| Kennebec | 1948 | US | 4n | Multi-purpose | 5061 | - | - | 4n | - |
| King Russet | 2018 | Netherlands | 4n | Fries processing | 5089 | - | - | 4n | - |
| Kolibri | 1998 | Germany | 4n | Crisps processing | 5202 | - | - | 4n | - |
| Krone | 2002 | Germany | 4n | Multi-purpose | 5220 | - | - | 4n | - |
| Kuba | 2005 | Poland | 4n | Starch processing | 5342 | - | - | 4n | - |
| Kuras | 1999 | Netherlands | 4n | Starch processing | 13-017- 3 | - | - | 4n | - |
| Lady Rosetta | 2004 | Netherlands | 4n | Chip processing | 14-214- 1 | - | - | 4n | - |
| Laura | 1988 | Germany | 4n | Salad making | 14-317- 1 | - | - | 4n | - |
| Leyla | 1988 | Germany | 4n | Multi-purpose | 14-912- 2 | - | - | 4n | - |

**Table S2: Genome-wide sequence variant distribution in 100 PotatoTools Cult21PT_950K clones**. Length (bp) is from the assembly of dAg1_v1.0 potato genome. The number of sequence variants are presented for individual chromosomes before and after quality control filtering (no monomorphic calls, Phred-quality score $\geq$ 20; average read depth of $\geq$15, depth (DP) lower than the average sequence DP divided by total clones plus three standard deviations, missing rate per polymorphism $\leq$20% and minor allele requency (MAF $>$ 0.05)). The average distance in bp's is given for the sequence variants after filtering.

| Chromosome | Length (bp) | Total sequence variants | Filtered sequence variants | Avg. distance (bp) |
|---|---|---|---|---|
| Chr01 | 89,719,359 | 7,464,200 | 2,848,988 | 35.51 |
| Chr02 | 52,458,522 | 4,532,819 | 1,382,111 | 37.95 |
| Chr03 | 61,672,818 | 4,915,225 | 1,419,247 | 43.45 |
| Chr04 | 73,201,088 | 6,341,880 | 1,968,319 | 37.18 |
| Chr05 | 56,731,602 | 5,024,190 | 1,836,972 | 30.88 |
| Chr06 | 55,985,673 | 4,194,361 | 1,381,789 | 40.51 |
| Chr07 | 53,532,588 | 4,969,769 | 1,571,783 | 34.05 |
| Chr08 | 63,071,485 | 5,488,133 | 1,654,632 | 38.11 |
| Chr09 | 73,873,122 | 6,516,524 | 1,886,027 | 39.16 |
| Chr10 | 57,411,799 | 3,724,762 | 1,128,971 | 50.85 |
| Chr11 | 55,350,225 | 4,949,674 | 1,579,120 | 35.05 |
| Chr12 | 57,049,570 | 5,190,007 | 1,478,389 | 38.58 |
| Chr_Unknown | 68,158,033 | 5,005,961 | 1,919,188 | 35.51 |
| Total | 818,215,884 | 68,317,505 | 22,055,536 | 37.90 |

**Table S3: Genome-wide raw and filtered sequence variant distribution in PotatoTools Non-Cult21PT_950K clones**. Sequence variants are presented for individual chromosomes before and after quality control filtering, and the average distance in bp's for filtered sequence variants and insertions/deletions (Indels) combined.

| **Clones** | **Raw sequence variants** | **Filtered SNPs** | **Filtered Indels** | **Avg. distance** |
|---|---|---|---|---|
| *S. hawkesianum* | 20,106,440 | 1,134,257 | 99,270 | 699.02 bp |
| *S. sparsipilum* | 24,615,589 | 1,800,813 | 166,243 | 417.11 bp |
| H80.577/1 | 16,923,925 | 1,159,782 | 123,856 | 659.31 bp |
| P40 | 22,847,580 | 1,626,684 | 180,377 | 464.16 bp |
| *S. gourlayi* | 21,513,265 | 1,483,461 | 131,918 | 528.44 bp |
| *S. spegazzini* | 20,777,208 | 928,523 | 83,208 | 818.90 bp |
| H98.D12 | 19,461,508 | 1,877,316 | 177,995 | 411.63 bp |
| *S. vernei* | 23,743,956 | 1,399,179 | 112,273 | 547.46 bp |
| Total | 169,989,471 | 11,410,015 | 1,075,140 | 568.25 bp |

**Table S4A: Number of unfiltered sequence variants located within varying distances in kilobase (Kbs) from potato genes**. Sequence variants were grouped into five categories: ≤1Kb, >1Kb and ≤2Kb, >2Kb and ≤3Kb, >3Kb and ≤4Kb, and >4Kb of gene.

| **Chromosome** | **≤1Kb** | **>1Kb and ≤2Kb** | **>2Kb and ≤3Kb** | **>3Kb and ≤4Kb** | **>4Kb** |
|---|---|---|---|---|---|
| Chr01 | 25,358 | 16,863 | 12,022 | 8,914 | 46,214 |
| Chr02 | 16,705 | 11,584 | 8,304 | 5,916 | 26,162 |
| Chr03 | 17,588 | 12,441 | 8,843 | 6,426 | 31,087 |
| Chr04 | 17,434 | 12,012 | 8,706 | 6,575 | 38,649 |
| Chr05 | 14,728 | 10,023 | 7,184 | 5,322 | 30,100 |
| Chr06 | 15,599 | 11,099 | 7,664 | 5,602 | 29,599 |
| Chr07 | 13,165 | 9,031 | 6,841 | 5,177 | 28,341 |
| Chr08 | 13,467 | 9,614 | 7,084 | 5,258 | 34,002 |
| Chr09 | 15,227 | 10,398 | 7,766 | 5,666 | 39,491 |
| Chr10 | 13,140 | 9,407 | 6,810 | 4,826 | 28,983 |
| Chr11 | 12,632 | 9,020 | 6,658 | 4,952 | 28,261 |
| Chr12 | 14,882 | 10,308 | 7,437 | 5,424 | 28,537 |
| **Total** | 189,925 (21.7%) | 131,800 (15%) | 95,319 (10.9%) | 70,058 (8%) | 389,426 (44.4%) |

**Table S4B: Number of filtered high quality sequence variants located within varying distances in kilobase (Kbs) from potato genes**. Sequence variants were grouped into five categories for fitted calls: ≤1Kb, >1Kb and ≤2Kb, >2Kb and ≤3Kb, >3Kb and ≤4Kb, and >4Kb of gene.

| **Chromosome** | **≤1Kb** | **>1Kb and ≤2Kb** | **>2Kb and ≤3Kb** | **>3Kb and ≤4Kb** | **>4Kb** |
|---|---|---|---|---|---|
| Chr01 | 7,556 | 5,062 | 3,557 | 2,492 | 8,312 |
| Chr02 | 5,913 | 4,094 | 2,780 | 1,837 | 5,270 |
| Chr03 | 6,039 | 4,335 | 2,908 | 1,880 | 6,401 |
| Chr04 | 4,530 | 3,118 | 2,264 | 1,682 | 5,883 |
| Chr05 | 4,061 | 2,761 | 1,870 | 1,333 | 4,677 |
| Chr06 | 4,929 | 3,604 | 2,382 | 1,642 | 5,744 |
| Chr07 | 4,067 | 2,800 | 2,119 | 1,521 | 4,759 |
| Chr08 | 3,825 | 2,692 | 1,935 | 1,274 | 5,508 |
| Chr09 | 4,035 | 2,777 | 2,024 | 1,404 | 6,238 |
| Chr10 | 3,657 | 2,572 | 1,795 | 1,160 | 4,954 |
| Chr11 | 3,243 | 2,270 | 1,796 | 1,192 | 4,686 |
| Chr12 | 3,732 | 2,528 | 1,889 | 1,283 | 3,965 |
| **Total** | 55,587 (27%) | 38,613 (18.7%) | 27,319 (13.2%) | 18,700(9.1%) | 66,397 (32.1%) |

**Table S5: Summary of fitted sequence variants obtained from fitPoly specific to Illumina Infinium 21K array (GGP3), PotatoTools Cult21PT_950K, and Non-Cult21PT_950K clones specific sequence variants per chromosome.**

| Chromosome | Total sequence variants | Fitted sequence variants | NonCult21PT_950K sequence variants | GGP3 and Cult21PT_950K sequence variants |
|---|---|---|---|---|
| Chr01 | 111,807 | 100,792 | 4,995 | 95,714 |
| Chr02 | 70,938 | 64,269 | 2,762 | 61,507 |
| Chr03 | 79,868 | 70,823 | 3,250 | 67,573 |
| Chr04 | 85,118 | 76,201 | 3,718 | 72,483 |
| Chr05 | 68,949 | 61,784 | 3,023 | 58,761 |
| Chr06 | 70,541 | 64,526 | 2,970 | 61,556 |
| Chr07 | 64,292 | 57,888 | 2,859 | 55,029 |
| Chr08 | 70,651 | 63,588 | 3,091 | 60,497 |
| Chr09 | 78,067 | 71,556 | 3,665 | 67,891 |
| Chr10 | 64,016 | 57,232 | 2,763 | 54,469 |
| Chr11 | 62,946 | 56,682 | 2,635 | 54,047 |
| Chr12 | 67,777 | 60,845 | 2,766 | 58,079 |
| Chr_Unknown | 52,600 | 46,548 | 2,780 | 43,768 |
| Chr00 | 128 | 118 | 0 | 118 |
| Chloroplast | 25 | 23 | 0 | 23 |
| Total | 947,723 | 852,792 | 41,277 | 811,515 |

**Table S6: Genome-wide distribution of genotyping calls filtered on minor allele frequency (MAF) and SNP call rate (CR) thresholds.** Illumina Infinium 21K array (GGP3), and PotatoTools Cult21PT_950K clone specific sequence variants were filtered on minor allele frequency (MAF) threshold of 0.01, SNP call rate (CR ≥80) and (CR ≥90).

| | Fitted calls | | MAF ≥ 0.01 | | MAF ≥ 0.01 and CR ≥ 80 | | MAF ≥ 0.01 and CR ≥ 90 | |
|---|---|---|---|---|---|---|---|---|
| **Chromosome** | **GGP3** | **Cult21PT_950K** | **GGP3** | **Cult21PT_950K** | **GGP3** | **Cult21PT_950K** | **GGP3** | **Cult21PT_950K** |
| Chr01 | 2,329 | 94,007 | 2,091 (89.8 %) | 85,825 (91.3%) | 1,982 (85.1%) | 68,378 (72.7%) | 1,850 (79.4%) | 54,600 (58.1%) |
| Chr02 | 1,792 | 60,075 | 1,577 (88%) | 55,534 (92.4%) | 1,469 (82%) | 45,341 (75.5%) | 1,375 (76.7%) | 36,814 (61.3%) |
| Chr03 | 1,627 | 66,379 | 1,487 (91.4%) | 60,882 (91.7%) | 1,409 (86.6%) | 49,442 (74.5%) | 1,319 (81.1%) | 40,647 (61.2%) |
| Chr04 | 1,665 | 71,287 | 1,493 (89.7%) | 64,720 (90.8%) | 1,398 (84%) | 50,076 (70.2%) | 1,295 (77.8%) | 38,785 (54.4%) |
| Chr05 | 1,516 | 57,666 | 1,420 (93.7%) | 52,457 (91%) | 1,334 (88%) | 40,798 (70.7%) | 1,235 (81.5%) | 31,386 (54.4%) |
| Chr06 | 1,325 | 60,609 | 1,249 (94.3%) | 55,629 (91.8%) | 1,175 (88.8%) | 44,923 (74.1%) | 1,086 (82%) | 36,638 (60.4%) |
| Chr07 | 1,411 | 53,994 | 1,245 (88.2%) | 49,454 (91.6%) | 1,167 (82.7%) | 39,207 (72.6%) | 1,080 (76.5%) | 30,792 (57%) |
| Chr08 | 1,187 | 59,734 | 1,073 (90.4%) | 54,608 (91.4%) | 1,012 (85.3%) | 42,806 (71.7%) | 926 (78%) | 33,750 (56.5%) |
| Chr09 | 1,263 | 67,105 | 1,171 (92.7%) | 60,916 (90.8%) | 1,090 (86.3%) | 47,817 (71.6%) | 995 (78.8%) | 37,636 (56.1%) |
| Chr10 | 1,068 | 53,769 | 949 (88.9%) | 49,003 (91.1%) | 887 (83.1%) | 39,384 (73.2%) | 821 (76.9%) | 32,287 (60%) |
| Chr11 | 1,359 | 53,065 | 1,251 (92.1%) | 48,051 (90.6%) | 1,169 (86%) | 37,615 (70.9%) | 1,053 (77.5%) | 29,322 (55.3%) |
| Chr12 | 1,124 | 57,372 | 1,007 (89.6%) | 52,064 (90.7%) | 954 (84.9%) | 40,671 (70.9%) | 875 (77.8 %) | 31,489 (54.9 %) |
| Chr_Unknown | 0 | 44,148 | 0 (0%) | 39,616 (89.7%) | 0 (0%) | 30,036 (68%) | 0 (0%) | 22,493 (50.9%) |
| Chloroplast | 118 | 0 | 115 (97.5%) | 0 (0%) | 108 (91.5%) | 0 (0%) | 95 (80.5%) | 0 (0%) |
| Chr00 | 23 | 0 | 18 (78.3%) | 0 (0%) | 16 (69.6%) | 0 (0%) | 16 (69.6%) | 0 (0%) |
| Total | 17,807 | 799,210 | 16,146 (90.7%) | 728,759 (91.2%) | 15,170 (85.2%) | 576,494 (72.1%) | 14,021 (78.7%) | 456,639 (57.1%) |

**Table S7: Genotype similarity for shared sequence variants between SolCap SNP array and PotatoTools Cult21PT_950K and NonCult21PT_950K clones specific sequence variants filtered on SNP call rate (CR≥ 90%) and minor allele frequency (MAF) ≥0.01**

| Clones | No. of sequence variants | Concordance % |
|---|---|---|
| Adretta | 5,367 | 94.2% |
| Agria_rep1 | 5,345 | 94.4% |
| Agria_rep2 | 5,367 | 94.5% |
| Agria_rep3 | 5,350 | 94.5% |
| Agria_rep4 | 5,323 | 94.4% |
| Agria_rep5 | 5,347 | 94.4 % |
| Agria_rep6 | 5,343 | 94.4% |
| Agria_rep7 | 5,355 | 94.5% |
| Agria_rep8 | 5,342 | 94.4% |
| Agria_rep9 | 5,354 | 94.5% |
| Agria_rep10 | 5,312 | 94.4% |
| Agria_rep11 | 5,350 | 94.4% |
| Agria_rep12 | 5,300 | 94.5% |
| Cara | 5,294 | 94.5% |
| Desiree | 5,325 | 94.4% |
| Flava | 5,319 | 94.4% |
| Gala | 5,336 | 94.5% |
| Granola | 5,314 | 94.3% |
| Innovator_rep1 | 5,327 | 94% |
| Innovator_rep2 | 5,350 | 94% |
| Kolibri | 5,356 | 94.3% |
| Krone | 5,373 | 94.2% |
| Kuba | 5,058 | 84% |
| Lady Rosetta | 5,330 | 94% |
| Marabel | 5,363 | 94.5% |
| Maris piper | 5,357 | 81.2% |
| Nicola | 5,369 | 94.3% |
| Quarta_rep1 | 5,352 | 94.2% |
| Quarta_rep2 | 5,357 | 80% |
| Schwalbe | 5,363 | 94.4% |
| **Average** | 5,333 | 93% |

**Table S8: Proportion of correct (AAAA, AABB, BBBB) and wrong allele calls (AAAB, ABBB) of diploids in a tetraploid genotyping space with SNP call rate ≥ 90% for Illumina Infinium 21K array (GGP3), PotatoTools NonCult21PT_950K and Cult21PT_950K clone specific sequence variants.** In a tetraploid genotyping space, the diploid clones are expected to show AAAA, AABB and BBBB genotypes. Genotypes other than these are wrong calls. The proportion of wrong and correct calls are given for both PotatoTools Cult21PT_950K and NonCult21PT_950K clones specific and Illumina Infinium 21K array (GGP3) variants with at least one genotype called for diploid clones (selected variants). The number of calls identified corresponds to the sequence variants that have been filtered on SNP call rate (CR≥ 90) and minor allele frequency (MAF ≥ 0.01) (SNPset-I), SNP call rate ≥ 90%, MAF ≥ 0.01, Euclidean distance ≥0.05 and HetSo=-0.1 (SNPset-II), SNP call rate ≥ 90%, MAF ≥ 0.01, Euclidean distance ≥0.075 and HetSo=-0.1 (SNPset-III), and SNP call rate ≥ 90%, MAF ≥ 0.01, Euclidean distance ≥0.01 and HetSo=-0.1 (SNPset-IV).

| Description | GGP3 | | | NonCult21PT_950K and Cult21PT_950K | | |
|---|---|---|---|---|---|---|
| | Total sequence variants | Correct calls | Wrong calls | Total sequence variants | Correct calls | Wrong calls |
| SNPset-I | 13,981 (99.7%) | 7,994 (57%) | 5,987 (42.8%) | 475,473 (99.2%) | 193,818 (40.7%) | 281,655 (59.2%) |
| SNPset-II | 12,474 (99.9%) | 7,631 (61.1%) | 4,843 (38.8%) | 316,414 (99.8%) | 160,196 (50.6%) | 156,218 (49.3%) |
| SNPset-III | 12,413 (99.9%) | 7,603 (61.2%) | 4,810 (38.7%) | 314,276 (99.8%) | 159,698 (50.8%) | 154,578 (49.1%) |
| SNPset-IV | 12,366(99.9%) | 7,585 (61.3%) | 4,781 (38.7%) | 312,425 (99.8%) | 159,273 (51%) | 153,152 (49%) |

**Table S9: Environments (i.e. year-location combinations) that were used in the experiment and their respective properties.**

| Environment | No. of entries | No. of populations | No. of blocks | No. of plants per plot |
|---|---|---|---|---|
| EUROPLANT 2019 Kaltenberg | 299 | 46 | 4 | 10 |
| EUROPLANT 2020 Kaltenberg | 297 | 47 | 4 | 16 |
| EUROPLANT 2020 Böhlendorf | 287 | 46 | 2 | 16 |
| EUROPLANT 2021 Kaltenberg | 300 | 48 | 4 | 16 |
| EUROPLANT 2021 Böhlendorf | 300 | 48 | 1 | 16 |
| Norika 2019 Groß Lüsewitz | 300 | 17 | 2 | 9 |
| Norika 2020 Groß Lüsewitz | 300 | 17 | 4 | 18 |
| Norika 2020 Mehringen | 300 | 17 | 3 | 20 |
| Norika 2021 Groß Lüsewitz | 297 | 17 | 4 | 18 |
| Norika 2021 Mehringen | 300 | 17 | 2 | 20 |
| Saka 2019 Windeby | 458 | 107 | 8 | 10 |
| Saka 2020 Windeby | 387 | 99 | 8 | 16 |
| Saka 2020 Gransebieth | 387 | 99 | 8 | 16 |
| Saka 2021 Windeby | 387 | 99 | 8 | 16 |
| Saka 2021 Gransebieth | 387 | 99 | 8 | 16 |

**Table S10: List of significant sequence variants along with their chromosomal coordinates and annotated genes identified in genome wide association study (GWAS) using the Potato Axiom 21PT_950K array to be associated with the polyphenol oxidase (PPO) trait.**

| Sequence variant | ID | Chromosome | Position | -log10(P-value) | Gene name |
|---|---|---|---|---|---|
| AX-611292391 | dAgria_v1.0_Chr01_176032 | Chr01 | 62543122 | 16.5 | NaN |
| AX-591421638 | dAgria_v1.0_Chr03_546095 | Chr03 | 10578483 | 13 | NaN |
| AX-608611998 | dAgria_v1.0_Chr08_1575286 | Chr08 | 6687777 | 7 | SOLTUB.AGRIA.G00000027135 |
| AX-592713689 | dAgria_v1.0_Chr08_1690697 | Chr08 | 51004690 | 9.8 | NaN |
| AX-592713987 | dAgria_v1.0_Chr08_1690959 | Chr08 | 51063330 | 14.6 | SOLTUB.AGRIA.G00000028381 |
| AX-592714022 | dAgria_v1.0_Chr08_1690985 | Chr08 | 51067372 | 6.6 | SOLTUB.AGRIA.G00000028382 |
| AX-592771820 | dAgria_v1.0_Chr08_1691468 | Chr08 | 51150513 | 13 | SOLTUB.AGRIA.G00000028392 |
| AX-592714578 | dAgria_v1.0_Chr08_1691487 | Chr08 | 51155041 | 22.1 | SOLTUB.AGRIA.G00000028392 |
| AX-592714689 | dAgria_v1.0_Chr08_1691578 | Chr08 | 51172497 | 9.3 | SOLTUB.AGRIA.G00000028393 |
| AX-592772352 | dAgria_v1.0_Chr08_1691943 | Chr08 | 51257786 | 11.2 | SOLTUB.AGRIA.G00000028402 |
| AX-592715145 | dAgria_v1.0_Chr08_1691959 | Chr08 | 51261300 | 11 | SOLTUB.AGRIA.G00000028402 |
| AX-592715150 | dAgria_v1.0_Chr08_1691966 | Chr08 | 51262007 | 13.6 | SOLTUB.AGRIA.G00000028402 |
| AX-592772384 | dAgria_v1.0_Chr08_1691974 | Chr08 | 51263260 | 13.1 | SOLTUB.AGRIA.G00000028402 |
| AX-592772474 | dAgria_v1.0_Chr08_1692043 | Chr08 | 51278522 | 15.4 | SOLTUB.AGRIA.G00000028403 |
| AX-592715349 | dAgria_v1.0_Chr08_1692135 | Chr08 | 51293882 | 8 | NaN |
| AX-592772605 | dAgria_v1.0_Chr08_1692167 | Chr08 | 51301134 | 6.8 | NaN |
| AX-608674565 | dAgria_v1.0_Chr08_1692325 | Chr08 | 51346304 | 34.8 | SOLTUB.AGRIA.G00000028409 |
| AX-592715563 | dAgria_v1.0_Chr08_1692328 | Chr08 | 51346588 | 16.2 | SOLTUB.AGRIA.G00000028409 |
| AX-608738588 | dAgria_v1.0_Chr08_1692357 | Chr08 | 51351550 | 13.7 | SOLTUB.AGRIA.G00000028409 |
| AX-592772856 | dAgria_v1.0_Chr08_1692373 | Chr08 | 51353341 | 9.6 | SOLTUB.AGRIA.G00000028409 |
| AX-608738605 | dAgria_v1.0_Chr08_1692377 | Chr08 | 51353816 | 35.2 | SOLTUB.AGRIA.G00000028409 |
| AX-592773051 | dAgria_v1.0_Chr08_1692545 | Chr08 | 51387159 | 28 | SOLTUB.AGRIA.G00000028413 |
| AX-608738817 | dAgria_v1.0_Chr08_1692597 | Chr08 | 51399593 | 30.9 | SOLTUB.AGRIA.G00000028413 |
| AX-592716019 | dAgria_v1.0_Chr08_1692716 | Chr08 | 51420985 | 16.6 | SOLTUB.AGRIA.G00000028415 |
| AX-608675004 | dAgria_v1.0_Chr08_1692769 | Chr08 | 51429193 | 15.9 | SOLTUB.AGRIA.G00000028416 |
| AX-608675444 | dAgria_v1.0_Chr08_1693239 | Chr08 | 51563487 | 8.3 | SOLTUB.AGRIA.G00000028424 |
| AX-592716883 | dAgria_v1.0_Chr08_1693440 | Chr08 | 51606971 | 36.4 | SOLTUB.AGRIA.G00000028429 |
| AX-612559949 | dAgria_v1.0_Chr08_1693465 | Chr08 | 51613651 | 39.6 | SOLTUB.AGRIA.G00000028429 |
| AX-592716994 | dAgria_v1.0_Chr08_1693533 | Chr08 | 51630584 | 14 | SOLTUB.AGRIA.G00000028429 |
| AX-592717071 | dAgria_v1.0_Chr08_1693582 | Chr08 | 51653956 | 16.7 | SOLTUB.AGRIA.G00000028429 |
| AX-592774614 | dAgria_v1.0_Chr08_1693861 | Chr08 | 51773549 | 8.8 | NaN |
| AX-612560465 | dAgria_v1.0_Chr08_1694432 | Chr08 | 51993597 | 45.2 | SOLTUB.AGRIA.G00000028441 |

| Sequence variant | ID | Chromosome | Position | -log10(P-value) | Gene name |
|---|---|---|---|---|---|
| AX-592718134 | dAgria_v1.0_Chr08_1694495 | Chr08 | 52,004,113 | 18.5 | SOLTUB.AGRIA.G00000028442 |
| AX-608741152 | dAgria_v1.0_Chr08_1694831 | Chr08 | 52,074,725 | 20.9 | SOLTUB.AGRIA.G00000028449 |
| AX-592775762 | dAgria_v1.0_Chr08_1694849 | Chr08 | 52,077,103 | 18.1 | SOLTUB.AGRIA.G00000028449 |
| AX-608677169 | dAgria_v1.0_Chr08_1694860 | Chr08 | 52,078,524 | 20.3 | SOLTUB.AGRIA.G00000028449 |
| AX-592718583 | dAgria_v1.0_Chr08_1694866 | Chr08 | 52,079,606 | 22.5 | SOLTUB.AGRIA.G00000028449 |
| AX-590715839 | dAgria_v1.0_H80_5771_Chr08_5907 | Chr08 | 52,296,819 | 12.9 | SOLTUB.AGRIA.G00000028464 |
| AX-592719664 | dAgria_v1.0_Chr08_1695804 | Chr08 | 52,314,835 | 18.7 | SOLTUB.AGRIA.G00000028466 |
| AX-606934136 | dAgria_v1.0_Chr08_1696062 | Chr08 | 52,378,012 | 13.4 | SOLTUB.AGRIA.G00000028469 |
| AX-608678437 | dAgria_v1.0_Chr08_1696165 | Chr08 | 52,393,488 | 18 | SOLTUB.AGRIA.G00000028471 |
| AX-592777293 | dAgria_v1.0_Chr08_1696220 | Chr08 | 52,407,646 | 16.6 | SOLTUB.AGRIA.G00000028472 |
| AX-592720203 | dAgria_v1.0_Chr08_1696309 | Chr08 | 52,427,050 | 19.1 | SOLTUB.AGRIA.G00000028474 |
| AX-592777416 | dAgria_v1.0_Chr08_1696312 | Chr08 | 52,427,737 | 15.5 | SOLTUB.AGRIA.G00000028474 |
| AX-592720548 | dAgria_v1.0_Chr08_1696598 | Chr08 | 52,514,471 | 7.5 | SOLTUB.AGRIA.G00000028477 |
| AX-592777853 | dAgria_v1.0_Chr08_1696706 | Chr08 | 52,533,163 | 14 | NaN |
| AX-592777881 | dAgria_v1.0_Chr08_1696726 | Chr08 | 52,536,898 | 13.3 | NaN |
| AX-592720694 | dAgria_v1.0_Chr08_1696732 | Chr08 | 52,538,086 | 19.9 | SOLTUB.AGRIA.G00000028483 |
| AX-592777917 | dAgria_v1.0_Chr08_1696750 | Chr08 | 52,540,623 | 18.8 | SOLTUB.AGRIA.G00000028483 |
| AX-592720811 | dAgria_v1.0_Chr08_1696837 | Chr08 | 52,557,328 | 21.7 | NaN |
| AX-608679133 | dAgria_v1.0_Chr08_1696838 | Chr08 | 52,557,367 | 22.2 | NaN |
| AX-608743382 | dAgria_v1.0_Chr08_1697059 | Chr08 | 52,615,748 | 18.2 | SOLTUB.AGRIA.G00000028489 |
| AX-592721074 | dAgria_v1.0_Chr08_1697062 | Chr08 | 52,616,380 | 14.8 | SOLTUB.AGRIA.G00000028489 |
| AX-592721274 | dAgria_v1.0_Chr08_1697237 | Chr08 | 52,665,801 | 20.8 | SOLTUB.AGRIA.G00000028491 |
| AX-592778616 | dAgria_v1.0_Chr08_1697360 | Chr08 | 52,697,770 | 16.5 | NaN |
| AX-606934269 | dAgria_v1.0_Chr08_1697416 | Chr08 | 52,703,823 | 21.8 | NaN |
| AX-592721529 | dAgria_v1.0_Chr08_1697490 | Chr08 | 52,710,242 | 46.7 | NaN |
| AX-592778894 | dAgria_v1.0_Chr08_1697611 | Chr08 | 52,742,338 | 27.5 | NaN |
| AX-634574794 | dAgria_v1.0_Chr08_3015947 | Chr08 | 52,926,760 | 15.1 | NaN |
| AX-608744655 | dAgria_v1.0_Chr08_1698363 | Chr08 | 52,943,988 | 6.9 | NaN |
| AX-592722641 | dAgria_v1.0_Chr08_1698393 | Chr08 | 52,959,761 | 9.4 | SOLTUB.AGRIA.G00000028513 |
| AX-592722708 | dAgria_v1.0_Chr08_1698445 | Chr08 | 52,971,511 | 11.4 | SOLTUB.AGRIA.G00000028513 |
| AX-592722765 | dAgria_v1.0_Chr08_1698497 | Chr08 | 52,981,872 | 21.3 | SOLTUB.AGRIA.G00000028516 |
| AX-612619969 | dAgria_v1.0_Chr08_1698619 | Chr08 | 53,019,138 | 9.5 | SOLTUB.AGRIA.G00000028517 |
| AX-608680842 | dAgria_v1.0_Chr08_1698621 | Chr08 | 53,019,232 | 9.3 | SOLTUB.AGRIA.G00000028517 |
| AX-608680844 | dAgria_v1.0_Chr08_1698622 | Chr08 | 53,020,113 | 19.4 | SOLTUB.AGRIA.G00000028517 |
| AX-608680882 | dAgria_v1.0_Chr08_1698670 | Chr08 | 53,039,849 | 18 | SOLTUB.AGRIA.G00000028517 |
| AX-593943825 | dAgria_v1.0_Chr08_1698673 | Chr08 | 53,040,910 | 17.8 | SOLTUB.AGRIA.G00000028517 |
| AX-608680884 | dAgria_v1.0_Chr08_1698676 | Chr08 | 53,041,194 | 18.6 | SOLTUB.AGRIA.G00000028517 |
| AX-608744947 | dAgria_v1.0_Chr08_1698677 | Chr08 | 53,041,645 | 18.1 | SOLTUB.AGRIA.G00000028517 |

| Sequence variant | ID | Chromosome | Position | -log10(P-value) | Gene name |
|---|---|---|---|---|---|
| AX-592723036 | dAgria_v1.0_Chr08_1698719 | Chr08 | 53,057,798 | 34.6 | SOLTUB.AGRIA.G00000028518 |
| AX-592780189 | dAgria_v1.0_Chr08_1698722 | Chr08 | 53,058,829 | 17.7 | SOLTUB.AGRIA.G00000028518 |
| AX-592723264 | dAgria_v1.0_Chr08_1698882 | Chr08 | 53,109,494 | 8.8 | SOLTUB.AGRIA.G00000028519 |
| AX-592780416 | dAgria_v1.0_Chr08_1698897 | Chr08 | 53,112,542 | 9.4 | SOLTUB.AGRIA.G00000028520 |
| AX-612620196 | dAgria_v1.0_Chr08_1699098 | Chr08 | 53,145,611 | 8.8 | SOLTUB.AGRIA.G00000028526 |
| AX-612562799 | dAgria_v1.0_Chr08_1699156 | Chr08 | 53,163,250 | 8.6 | SOLTUB.AGRIA.G00000028526 |
| AX-592780710 | dAgria_v1.0_Chr08_1699168 | Chr08 | 53,170,634 | 8.6 | SOLTUB.AGRIA.G00000028526 |
| AX-608681397 | dAgria_v1.0_Chr08_1699197 | Chr08 | 53,176,822 | 10.8 | SOLTUB.AGRIA.G00000028526 |
| AX-592724146 | dAgria_v1.0_Chr08_1699668 | Chr08 | 53,260,837 | 11.6 | SOLTUB.AGRIA.G00000028536 |
| AX-590642064 | dAgria_v1.0_H80$_5$771_$Chr$08_5921 | Chr08 | 53,303,788 | 9.7 | NaN |
| AX-592724868 | dAgria_v1.0_Chr08_1700285 | Chr08 | 53,383,813 | 6.6 | SOLTUB.AGRIA.G00000028548 |
| AX-635060907 | dAgria_v1.0_Chr08_3016177 | Chr08 | 53,953,052 | 7.3 | NaN |
| AX-592725735 | dAgria_v1.0_Chr08_1704127 | Chr08 | 54,180,967 | 7.9 | SOLTUB.AGRIA.G00000028633 |
| AX-612622539 | dAgria_v1.0_Chr08_1704152 | Chr08 | 54,185,825 | 7.2 | SOLTUB.AGRIA.G00000028633 |

**Table S11: List of significant sequence variants along with their chromosomal coordinates and annotated genes identified in genome wide association study (GWAS) using the Illumina Infinium 21K array (GGP3) specific variants linked with polyphenol oxidase (PPO).**

| Sequence variant | ID | Chromosome | Position | -log10(P-value) | Gene name |
|---|---|---|---|---|---|
| AX-593944261 | PotVar0077225 | Chr08 | 51,136,768 | 12 | SOLTUB.AGRIA.G00000028389, SOLTUB.AGRIA.G00000028390 |
| AX-593966525 | solcap_snp_c2_15803 | Chr08 | 51,421,811 | 22.4 | SOLTUB.AGRIA.G00000028415 |
| AX-593923809 | solcap_snp_c2_50702 | Chr08 | 51,985,513 | 47 | NaN |
| AX-593910856 | PotVar0125622 | Chr08 | 52,003,170 | 14 | SOLTUB.AGRIA.G00000028442 |

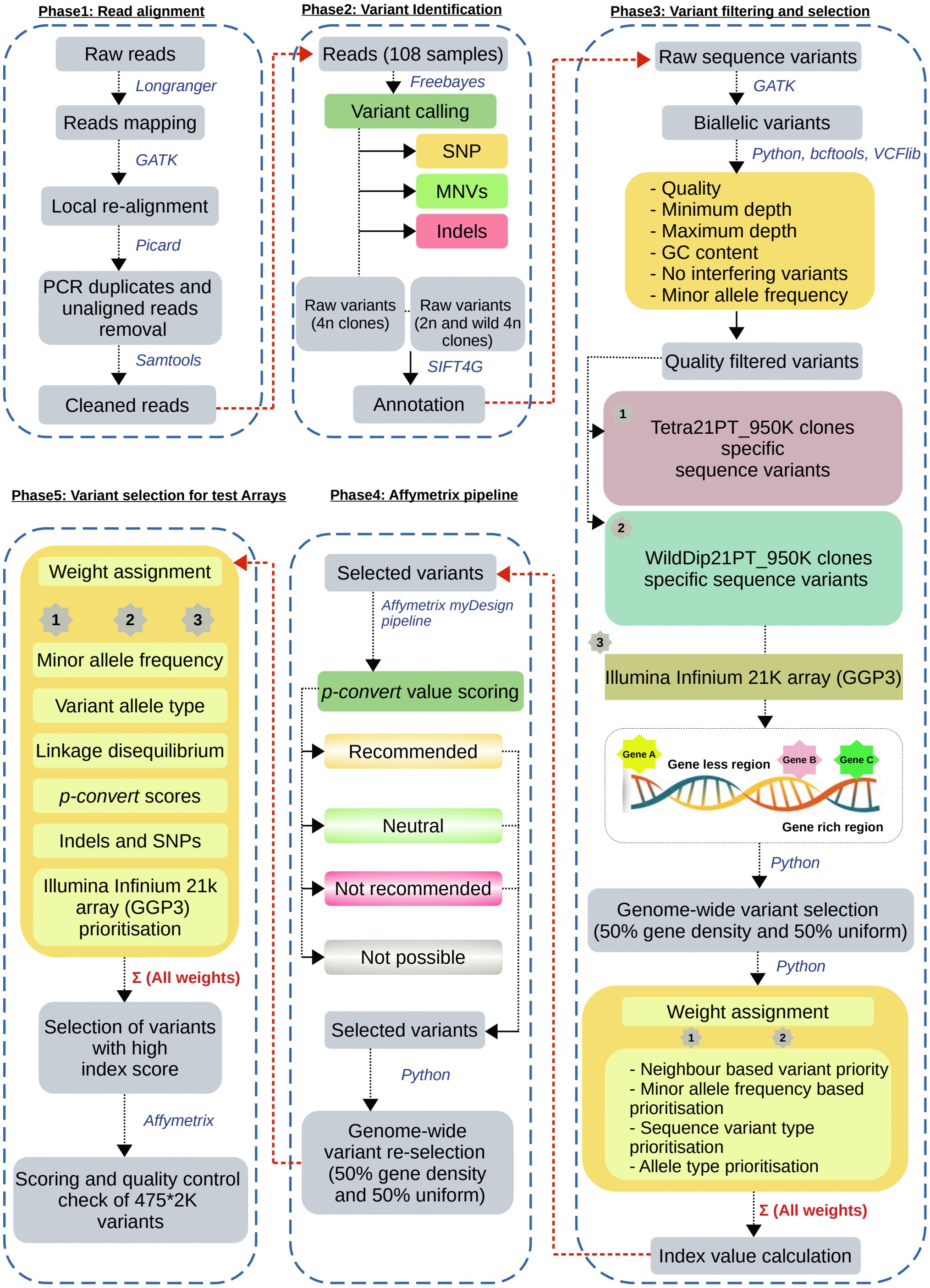


**Figure S1: Schematic workflow describing the development and evaluation of the Potato Axiom 21PT_950K screening arrays.** Workflow included 5 major phases (i) raw read alignment, (ii) sequence variant identification, (iii) variant filtering and selection (iv) sequence variant analysis by Affymetrix pipeline, and (v) final selection of sequence variants for tiling on the Potato Axiom 21PT_950K screening array.

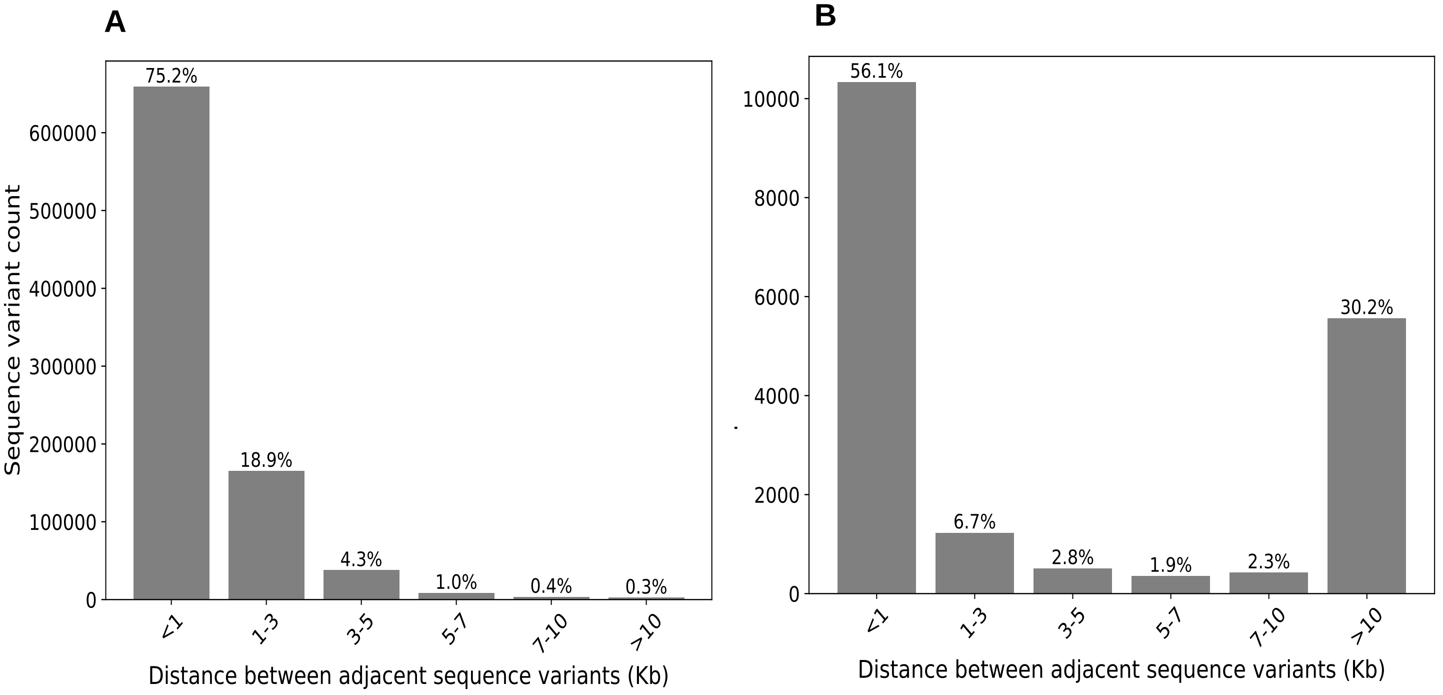


**Figure S2: Distribution of distances between sequence variants tiled on Potato Axiom 21PT_950K screening arrays design.** (A) Frequency distribution of adjacent inter-variant distances for PotatoTools Cult21PT_950K and NonCult21PT_950K clones specific sequence variants. The x-axis represents the frequency while the y-axis represents the distance between consecutive variants. (B) Frequency distribution for the distance in kilobase (Kb) between adjacent Illumina Infinium 21K array (GGP3) sequence variants.

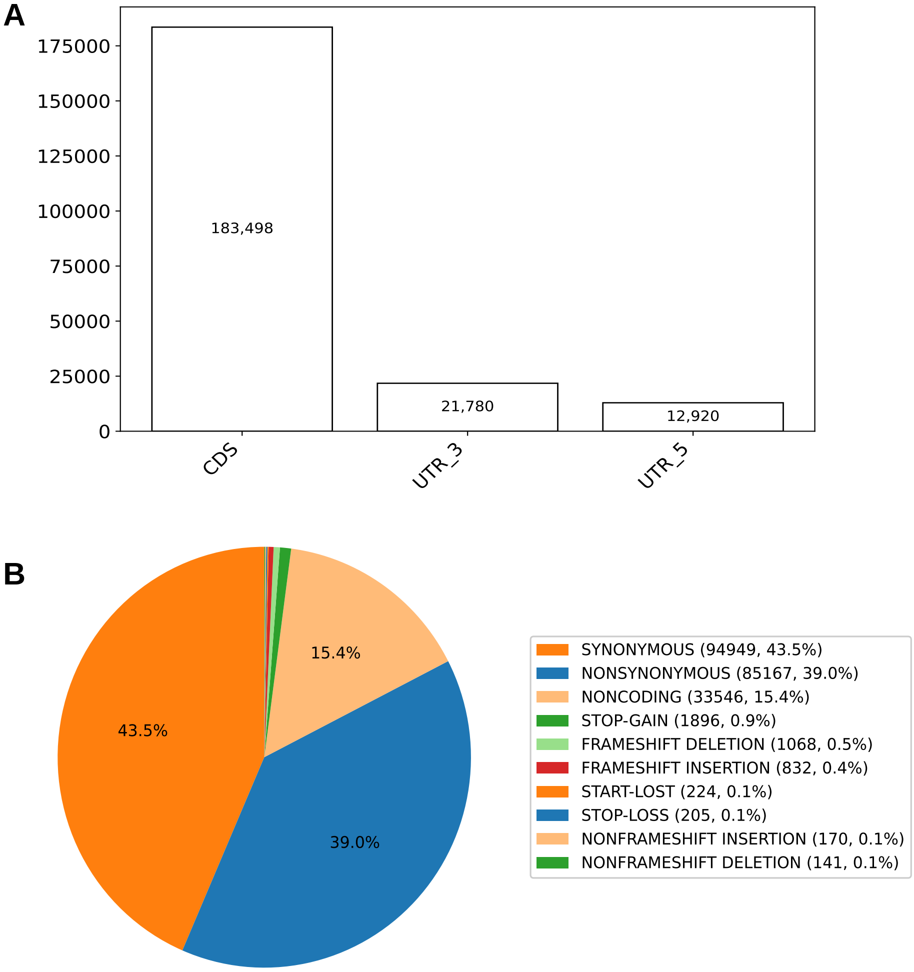


**Figure S3: SIFT4G based annotation of sequence variants** (A) Proportion of the sequence variants tiled on the potato Axiom_21PT_950K screening arrays annotated in coding sequence (CDS), three prime untranslated region (3′-UTR) and five prime untranslated region (5′-UTR). (B) Pie-chart representing the functional anotation of sequence variants based on SIFT4G.

A

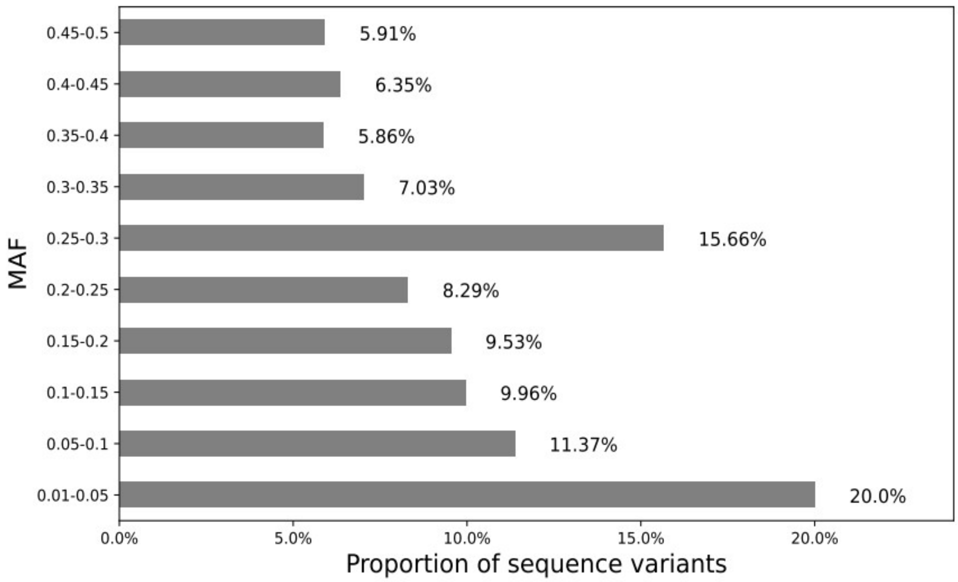


B

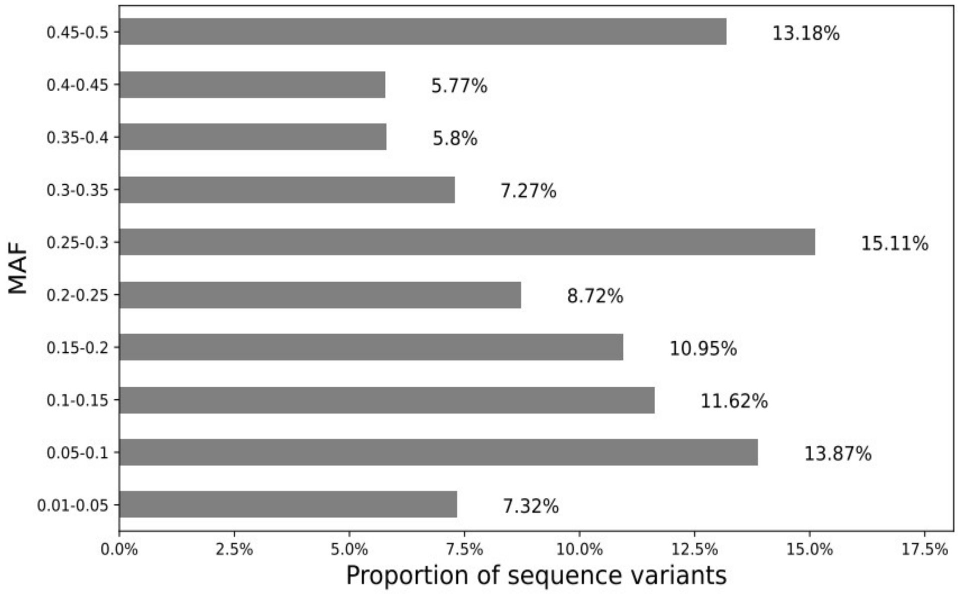


**Figure S4: Minor allele frequency (MAF) distribution in (A) PotatoTools Cult21PT_950K clones specific sequence variants and (B) Illumina Infinium 21K array (GGP3) filtered on SNP call rate (CR $\geq$ 90) and minor allele frequency (MAF $\geq$ 0.01).** The x-axis represents the proportion of sequence variants in percentage and y-axis represents the minor allele frequency (MAF) distribution.
